# Using spatio-temporal information in weather radar data to detect and track communal bird roosts

**DOI:** 10.1101/2022.10.28.513761

**Authors:** Gustavo Perez, Wenlong Zhao, Zezhou Cheng, Maria Carolina T. D. Belotti, Yuting Deng, Victoria F. Simons, Elske Tielens, Jeffrey F. Kelly, Kyle G. Horton, Subhransu Maji, Daniel Sheldon

**Author notes:** Equal contribution (Gustavo Perez, Wenlong Zhao).

## Abstract

1. The exodus of swallows from communal nighttime roosts is often visible as an expanding ring-shaped pattern in weather radar data. The WSR-88D network operated by the National Weather Service archives more than 25 years of data across 143 stations in the contiguous US. However, access to information about the roosting behavior of swallows is limited by the cost of manual annotation of these scans.
2. We develop an AI system to detect and track swallow roosts in weather radar data. Our model is based on the Faster R-CNN architecture and is customized to incorporate multiple spatial and temporal channels in volumetric radar scans using novel adaptor layers. We systematically study the impact of network architecture and input representation for this task. We incorporate our detection outputs into an AI-assisted system with an interface for human screening to collect research-grade data about roosting behavior. We deploy the system to collect information from 12 radar stations in the Great Lakes region of the US spanning 21 years.
3. The addition of temporal information improves roost detection performance from 47.0% mean average precision to 54.7%. Temporal information helps the model recognize the expanding pattern of roosts and filter false positives due to rain and static structures. Our system allowed the annotation of 15,628 roost signatures with 64,620 single-frame detections in 612,786 radar scans with 183.6 total hours of human screening, or 1.08 seconds per radar scan.
4. Our AI-assisted system provides research-quality roost data with far less human effort than manual annotation of radar scans. The data contains critical information about the phenology and population trends of swallows and martins, a declining group of aerial insectivores. Our successful deployment to collect historical data for 8% of the radar stations in the contiguous US lays the groundwork for continent-scale analysis of swallow roosts, and provides a starting point for analysis of other family-specific phenomena in weather radar, such as bat roosts and mayfly hatches.

## 1 Introduction

Monitoring animal populations is a cornerstone of ecological science. Information over large regions, long time periods, and with fine spatial and temporal resolution is increasingly important to understand the rapid and heterogeneous changes in animal populations due to climate change and human disturbance (Bairlein, 2016; Rosenberg *et al*., 2019; Sánderson *et al*., 2006; Sánchez-Bayo & Wyckhuys, 2019). Traditionally, gathering detailed and comprehensive population data has been extremely difficult, but technological advances can increase the scope and resolution of data by conducting monitoring passively, with less human effort, or by harnessing the power of large numbers of volunteers. Examples include digital tracking technologies (McKinnon & Love, 2018), camera traps (Burton *et al*., 2015), acoustic monitoring (Sugai *et al*., 2019; Gibb *et al*., 2019), and citizen science projects (Silvertown, 2009; Sullivan *et al*., 2014).

Weather radar is one of the most promising technologies for studying flying animals, with networks of weather surveillance radars around the globe continuously monitoring the airspace and detecting birds, bats, and insects in addition to precipitation (Kunz *et al*., 2008). The US NEXRAD weather radar network (Crum & Alberty, 1993), in particular, has archived 25 years of data from 143 radar stations covering nearly the entire contiguous US (Ansari *et al*., 2018), and offers the possibility of monitoring flying animals at an unprecedented scale and resolution (Gauthreaux, 1970; Bruderer, 1997; Gauthreaux & Belser, 1998; Gauthreaux *et al*., 2003; Dokter *et al*., 2011). These data have fueled a growing number of studies at increasing scales about bird populations, including studies about nocturnal migration and stopover behaviors (Gauthreaux *et al*., 2003; Farnsworth *et al*., 2016; Buler & Diehl, 2009; Buler & Dawson, 2014; Horton *et al*., 2018; Cohen *et al*., 2021), demography (Dokter *et al*., 2018b), the effects of artificial light on migration (Van Doren *et al*., 2017; McLaren *et al*., 2018; Van Doren *et al*., 2021), systems to forecast migration (Van Doren & Horton, 2018), and landmark findings about the declines (Rosenberg *et al*., 2019) and shifting phenologies of North American birds (Horton *et al*., 2020). NEXRAD data have also produced insights into changing bat (Stepanian & Wainwright, 2018) and insect (Stepanian *et al*., 2020) populations in North America, while similar research programs are being carried out on other continents (Shamoun-Baranes *et al*., 2014; Nilsson *et al*., 2019; Nussbaumer *et al*., 2019).

Algorithmic advances, including the use of artificial intelligence (AI), are essential for unlocking biological information contained in NEXRAD radar data. The archive includes more than 0.5 petabytes of data and 240 million scans, each of which may contain a variety of patterns corresponding to different types of precipitation, clutter, or biological scatterers. Methods are needed to automatically recognize, discriminate, and track different types of biological scatterers to collect measurements at large scales. AI methods based on deep learning and convolutional neural networks have shown tremendous success at related visual recognition tasks, and are excellent candidates for recognizing biological patterns in weather radar. Past work has focused on discriminating precipitation from broad-scale bird migration in radar data through the use of AI classification (RoyChowdhury *et al*., 2016; Van Doren & Horton, 2018; Horton *et al*., 2019b) or segmentation (Lin *et al*., 2019) methods.

This paper investigates an AI system to automatically detect and track martins and swallow roosts in NEXRAD radar data. Unlike broad-scale bird migration, swallow roosts are one of a relative small number radar phenomena that can be traced to family level.

A distinctive expanding ring pattern appears on radar as swallows and martins leave communal nocturnal roosting locations, ascend to elevations detectable by radar, and then disperse in different directions (Russell & Gauthreaux Jr, 1998) (also see Figure 1a and Figure 2). Communal roosts occur throughout the US, especially in late summer and fall (Russell *et al*., 1998), and may contain any of the seven North American swallow species (Winkler, 2006), though are often dominated by a single species, especially Purple Martins (*Progne subis*) and Tree Swallows (*Tachycineta bicolor*) (Bridge *et al*., 2016; Laughlin *et al*., 2016), depending on habitat and time of year. Swallows and martins are aerial insectivores, which are rapidly declining in North America (Nebel *et al*., 2010; Fraser *et al*., 2012; Rosenberg *et al*., 2019), so taxon-specific results about their ecology are of great interest.

**Figure 1:**
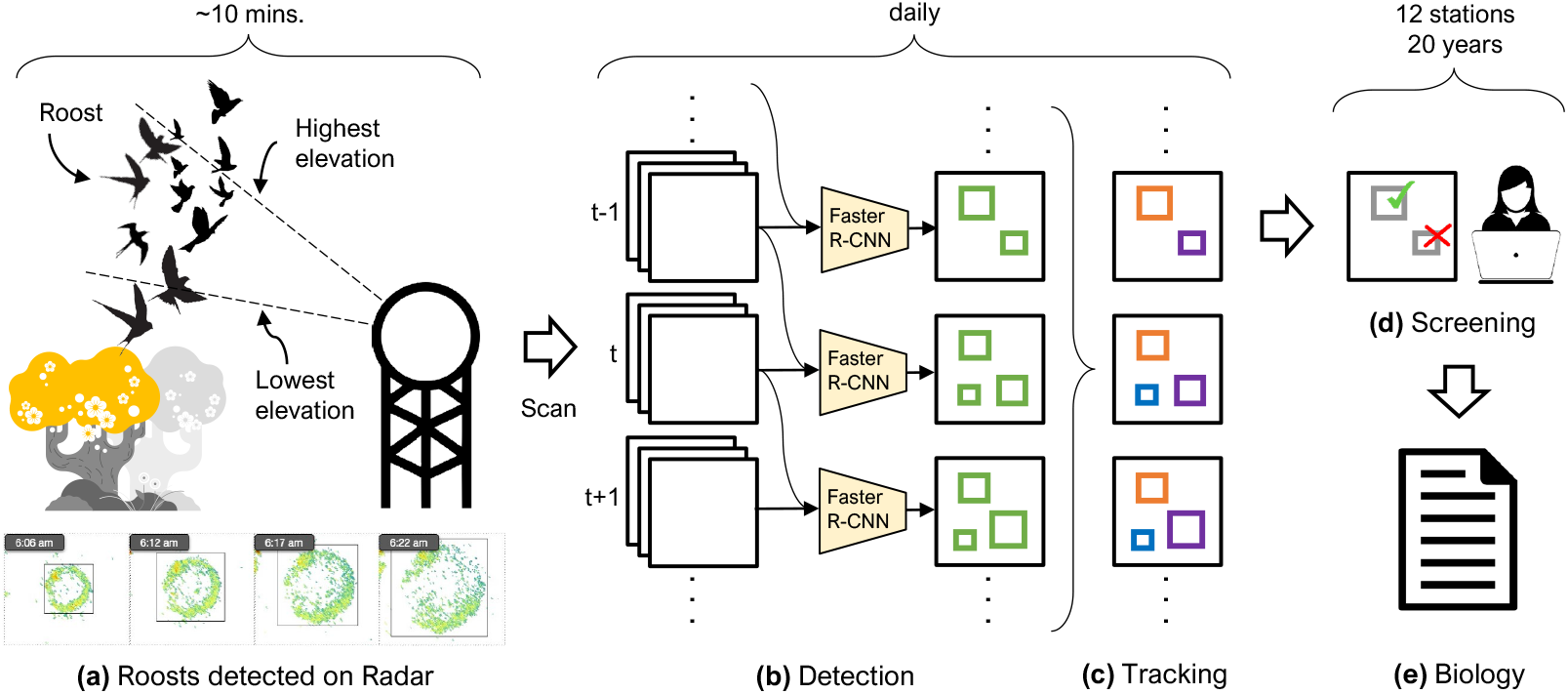
The proposed AI-assisted system. (a) The exodus of birds (e.g., swallows and martins) from nightly roosting locations are visible on weather radars as expanding rings in consecutive radar scans, particularly at the lowest elevations. (b-c) Our system detects individual roosts in each scan (shown as green boxes) using a spatio-temporal object detection model and organizes these into tracks corresponding to individual roosts every day (boxes of the same color correspond to the same track). (d) We have developed a web-based interface to enable quick screening of tracks, allowing us to analyze 20 years of data across 12 radar stations. (e) The screened data can provide insights about population trends and phenology of the tracked birds.

**Figure 2:**
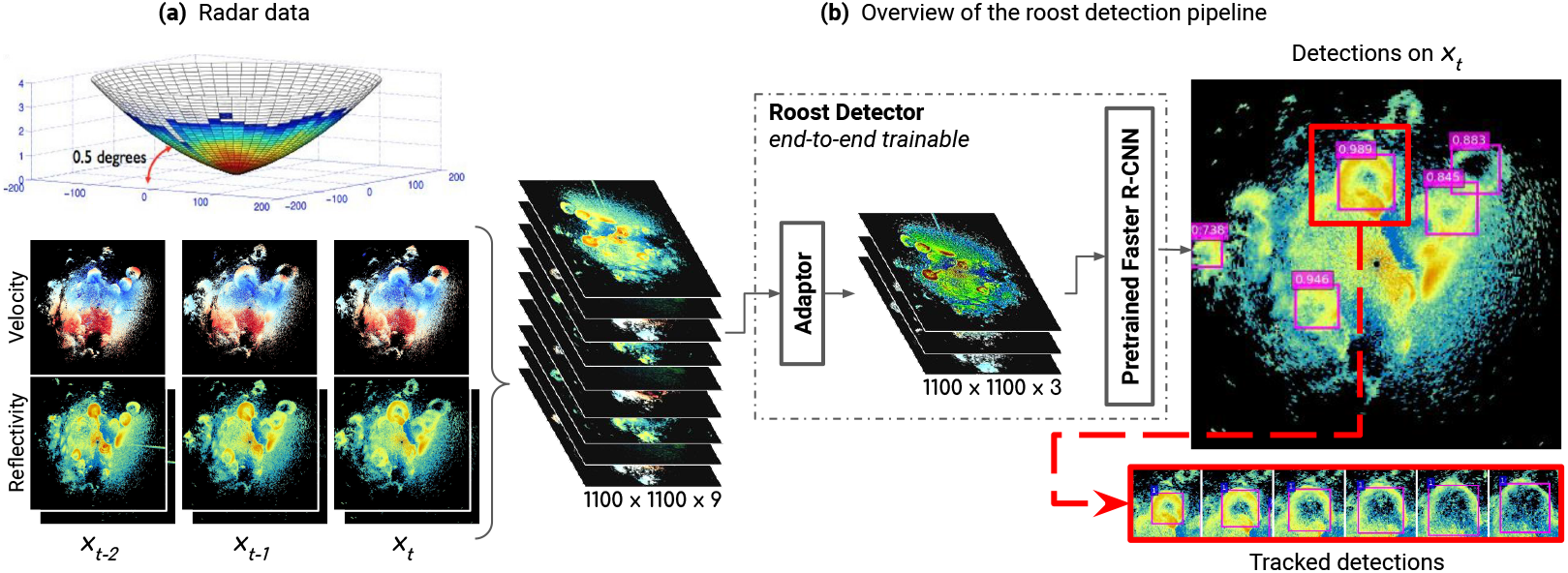
(a) Radar data. We show here a sweep at an elevation of 0.5°. We render top-down view of reflectivity channels at 0.5° and 1.5° and the velocity channel at 0.5° from 3 consecutive time frames *x*_*t−*2_, *x*_*t−*1_, and *x*_*t*_. **(b) Overview of the roost detection pipeline**. Radar images are concatenated into a single 9-channel image. The adaptor maps 9 channels to 3 channels to match a conventional RGB image for input into a deep network pre-trained on color images. A Faster R-CNN is trained to detect communal bird roosts for frame *x*_*t*_. Finally, we associate detections from individual frames into tracks.

An automated system to monitor martins and swallow roosting activity in past, present, and future radar scans could provide information urgently needed for basic science and conservation of these species. Past studies have used radar to study swallow roosts using manual annotation by humans to identify roosts in radar data (Winkler, 2006; Laughlin *et al*., 2013, 2016; Bridge *et al*., 2016; Kelly & Pletschet, 2017), but cannot be easily repeated or expanded due to the cost of human annotation. Chilson *et al.* (2018) trained a classifier to identify radar scans that contained roosts. From an AI perspective, roost detection is most naturally posed as an object detection and tracking problem. We previously developed the first AI system for detecting and tracking roosts (Cheng *et al*., 2020). That work focused on training procedures to deal with inconsistent annotation styles, and the resulting system detected obvious roosts with few mistakes, but included a number of false positives from rain and other sources when tuned to detect a large fraction of the human-annotated roosts. Thus, it could not fully automatically generate high-quality data for research purposes.

In this paper, we investigate the design, deployment, and analysis of an AI-assisted system capable of extracting research-grade roost data from NEXRAD radar scans as shown in Figure 1. Building on our prior system, we make a number of novel contributions. We describe the computer vision background and methods in a way accessible to biology researchers and provide open source implementations of our training procedures and deployed models. We construct a standardized dataset of roost annotations to remove labeling style differences and to support standard training and evaluation methods. We augment the neural network architecture to accept additional channels of input data, including radar moments at different elevations as well as from previous and future scans, which allows the network to recognize the distinctive expanding movement dynamics of roosts rings. The use of temporal information and other model improvements lead to increased performance compared to Cheng *et al.* (2020). We report on the deployment of the system to extract 21 years of data for 12 radar stations in the Great Lakes region of the US, for which we developed a software system and screening protocol for our team to screen the AI model’s outputs for errors. With this AI-assisted system, we successfully extracted research-grade historical roost data for 8% of the radar stations in the contiguous US covering a region with large swallow populations. We demonstrate the types of biological research possible with this data through a case study of historical swallow populations on Walpole Island in Ontario, Canada.

## 2 Materials and Methods

We aim to develop a machine learning system that automatically detects and tracks communally roosting birds in weather radar data, thus overcoming the intractability of manual annotations and producing high-quality data for radar aeroecology research.

### 2.1 Radar Data Preliminaries

#### Radar Data

The Weather Surveillance Radar-1988 Doppler (WSR-88D, also called NEXRAD) network operated by the U.S. National Weather Service contains 143 stations in the contiguous U.S. and another 16 stations in Alaska, Hawaii, and other U.S. territories. It has archived raster-like radar data products since the 1990s. The data products are collected through radar volume scans, in which the radar antenna rotates 360° around the vertical axis to sample cone-shaped “slices” of the surrounding airspace at different elevation angles (Figure 2a top). The data are collected every 4-10 minutes.

Conventionally, WSR-88D radars collect 3 radar moments at various elevation angles. *Reflectivity factor* measures the density of objects in the atmosphere. Radial velocity measures the speed at which objects are moving relative to the radar station using the Doppler shift of the reflected radio waves. Reflectivity-weighted mean and standard deviation of radial velocity are collected as the *radial velocity* and *spectrum width* radar moments. In this paper, our detectors are trained using subsets of these products or “channels”. Between 2011 and 2013, the radar stations were upgraded to also collect three dual polarization (“dual-pol”) radar moments (Stepanian, 2015; Stepanian *et al*., 2016): *differential reflectivity, differential phase*, and *correlation coefficient*. These products result from both horizontal and vertical radar waves, and can effectively been used to identify rain (Zrnić & Ryzhkov, 1998; Stepanian *et al*., 2016; Dokter *et al*., 2018a; Cheng *et al*., 2020). We use dual-pol products, whenever available, to reduce false roost detections caused by rain during deployment of our system.

#### Rendering

WSR-88D radar sweeps produce two-dimensional arrays in polar coordinates indexed by range and azimuth (antenna pointing direction in the horizontal plane), with fixed antenna elevation angle. One volume scan consists of sweeps for each radar product for a sequence of increasing elevation angles. We use nearest neighbor interpolation (Parker *et al*., 1983) to resample each sweep onto a fixed 600 × 600 Cartesian grid centered at the radar station with 500 m pixels out to a radius of 150 km. This rendering is known as a *plan position indicator* and corresponds to a top-down view of the cone shown at the top of Figure 2a. For each product in each volume scan, we produce a threedimensional grid. The first dimension is indexed by the five elevation angles 0.5°, 1.5°, 2.5°, 3.5°, and 4.5°, while the second and third dimensions correspond to the 600 × 600 Cartesian grid. To populate the array, we choose the five sweeps with elevation angles closest to the ones indicated above; scanning patterns are variable but almost always include sweeps at or near these five elevation angles.

### 2.2 Object Detection and Tracking Preliminaries

It is natural to frame the problem of identifying roost signatures in radar data as an object detection and tracking problem. In this section we review the computer vision techniques needed to approach these tasks and adapt them to roost detection and tracking.

#### Object detection

Object detection aims to identify the category and location of objects in images represented by their circumscribing box (or bounding box). The basic approach for object detection is to apply a classifier (e.g., roost or not) at multiple locations within the image followed by a procedure to aggregate scores from all locations to produce a non-overlapping set of detections, which is often referred to as “non-maximum suppression”. A challenge is that the process is computationally demanding: one has to search over a large number of locations and scales corresponding to different positions, size and aspect ratios of the objects. Early detectors mitigated this by applying simple classifiers to image patches at densely sampled locations in an image (Viola & Jones, 2001; Dalal & Triggs, 2005), or expensive classifiers based on complex features on a small set of locations obtained using a proposal generation mechanism (Girshick *et al*., 2016; Malisiewicz *et al*., 2011; Chum & Zisserman, 2007). In either case, image patches are represented as a set of features suitable for a machine learning classifier. Breakthroughs in deep learning, in particular advances in training deep convolutional neural networks (CNNs) on large image datasets such as ImageNet (Deng *et al*., 2009) have largely replaced the hand-designed features and have produced significant improvements on various object detection benchmarks. Our earlier work (Cheng *et al*., 2020) found that detectors based on Faster R-CNNs (Ren *et al*., 2015) were effective at detecting roosts, and forms the basis of this work.

Faster R-CNN combines a region proposal network (RPN) with a detection network to predict the category scores and bounding-boxes of each proposal. The RPN and detection network share most of the computations by building on top of a shared feature extraction network (“backbone”), which results in a faster processing time compared to earlier designs. For example, R-CNN (Girshick *et al*., 2016) used classical region proposals based on color and texture and applied the CNN classifier to each of these proposals, while Fast R-CNN (Girshick, 2015) improved the processing time by re-sampling pre-computed CNN features on the proposals via a region-of-interest (ROI) pooling layer instead of processing them independently. Faster R-CNN combines all these steps (described in detail below and Figure 3) resulting in a single end-to-end trainable model.

**Figure 3:**
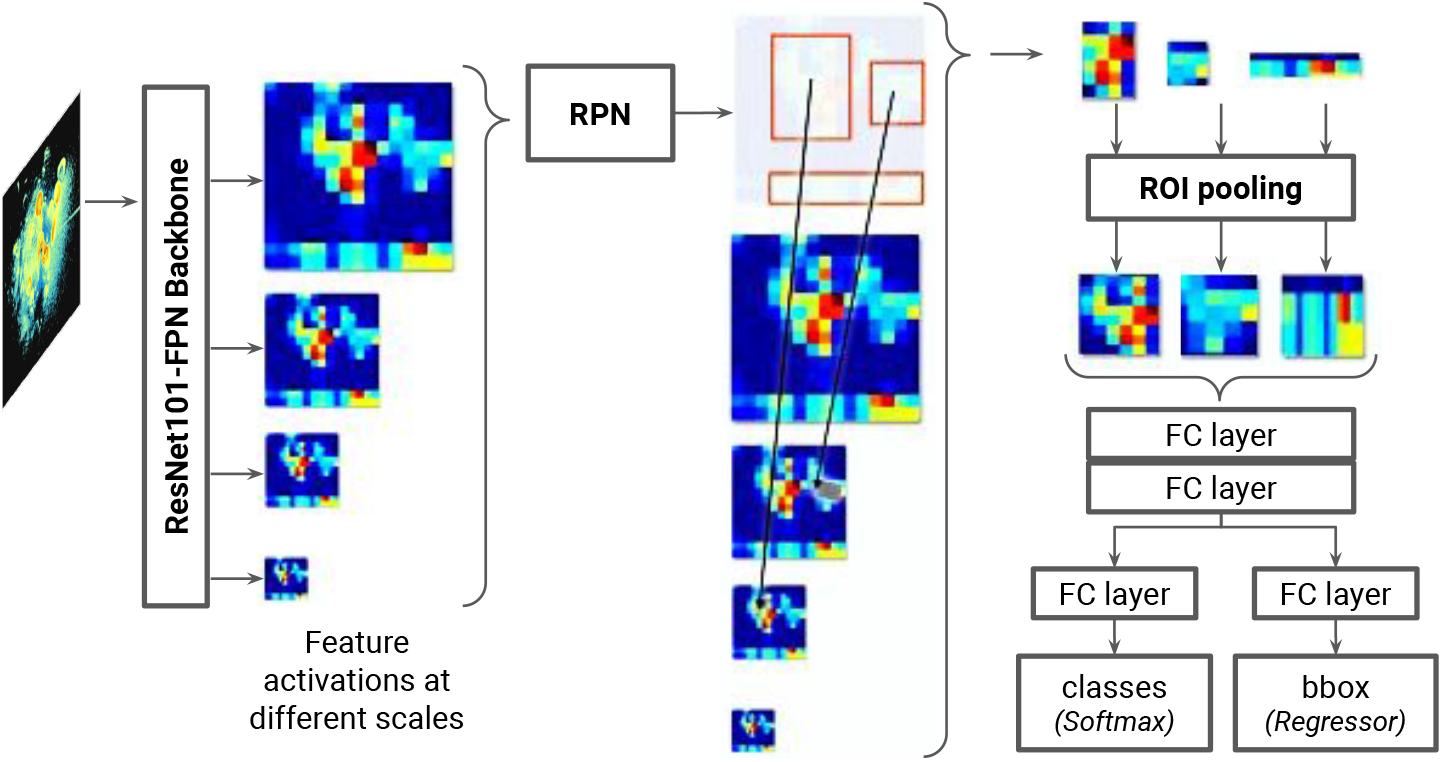
A Faster R-CNN based roost detector on radar images. The detector uses a feature pyramid network (FPN) to extract multi-scale features from the input. The region proposal network (RPN) uses these to generate object proposals across scales and locations which indicate potential regions that contain roosts. The features are re-sampled at the proposals by a region-of-interest (ROI) pooling layer to obtain a fixed dimensional representation of each proposal. Finally, these are classified as roosts or background, and the bounding-box is re-estimated.

- Region proposal network (RPN): The RPN ranks “anchors” (region bounding boxes) and proposes those with a higher likelihood of containing an object. It predicts a probability of an anchor to be an object or background, and it refines each anchor’s size with a bounding box regressor.
- Anchors: An anchor is a box with a fixed pre-defined size and height-width ratio. Faster R-CNN’s default configuration uses 15 anchors at each position in the image: 5 sizes (32, 64, 128, 256, 512)^1^ at three different ratios (1:1, 1:2, and 2:1). The anchors’ sizes are hyper-parameters and they will affect the performance of the object detector depending on each application. In radar images, roosts are mostly isometric and appear from early stages until they are completely dissolved, so we use a single ratio of 1:1 with a larger range of anchor sizes.
- ROI pooling: The RPN outputs region proposals of different sizes. To use the feature maps of each proposed region, the region of interest pooling (ROI pooling) reduces the features to the same size by dividing the region into a fixed number of approximately equally sized regions and then applies *max* pooling to each one.

#### Convolutional Neural Networks (CNNs)

CNNs are deep networks designed for image processing. They consist of operations organized in “layers” of computation including simple functions such as convolutions, entrywise non-linear transformations, spatial aggregation within regions (or “pooling”), and others such as normalization, linear transformations, and more. These operations use learned network parameters to map the input (e.g., an image) to the output (e.g., a classification score). While early CNNs (LeCun *et al*., 1998) were small and trained on a simple task such as digit and alphabet recognition, modern CNNs trained on large-scale datasets such as ImageNet and MS-COCO (Lin *et al*., 2014) contain tens of millions of parameters. Furthermore, the “activations”, or outputs of intermediate layers, extracted from CNNs pre-trained on ImageNet have been shown to contain rich, hierarchical representations suitable for many image understanding tasks such as object categorization, shape understanding, and texture recognition.

We use a Residual Network (ResNet) “backbone” (He *et al*., 2015) for our object detector. This network is the basis of many state-of-the-art detectors in computer vision, and has shown better generalization and training stability than some of the earlier networks such as AlexNet (Krizhevsky *et al*., 2012) and VGGNet (Simonyan & Zisserman, 2015). In particular we use ResNet50-FPN and ResNet101-FPN, corresponding to 50 and 101 layers respectively. These networks extract multi-scale features of the input organized as a feature pyramid network (FPN) allowing better detection of objects across scales (See Figure 3). The RPN and detection networks described in the earlier section are attached to the end of the ResNet backbone to train the Faster R-CNN model. See the book of (Goodfellow *et al*., 2016) for a detailed background on CNNs.

#### Adaptors

Many studies (Yosinski *et al*., 2014; Zhuang *et al*., 2019; Ribani & Marengoni, 2019) have shown that pre-trained networks lead to faster convergence and greater robustness to hyperparameter settings when training object detectors. However, to use networks pre-trained on three-channel (RGB) images we must adapt the input to account for the different number of channels available for radar volume scans, which include the different radar products, elevation angles, and potentially sweeps from temporally-adjacent scans. We propose a learnable adaptor, which maps an arbitrary *k*-channel image *x* ∈ℝ^*k×n×m*^ to a 3-channel image using a linear projection. This is implemented as a single convolutional layer with three filters of size *k* ×1 × 1, each of which implements a (learned) linear mapping from ℝ^*k*^ to ℝ^1^. While other choices are possible, including a non-linear adaptor, or replicating filters in the first layer of the network to match the shape of an input with more channels, our preliminary study suggested that linear adaptors are the most effective. See (Perez & Maji, 2022) for a study on how the architecture of adaptor affects transfer learning. In Section 3.2 and Table 3b we evaluate the benefits of pre-training.

#### Training and Evaluation

The parameters *θ* of the model, which consist of all learnable weights of the neural network components shown in Figure 3, are learned by minimizing a multi-task loss function *L* = *L*_*cls*_ + *λL*_*reg*_, where *L*_*cls*_ is a classification loss (log-likeilhood of the binary logistic model for “roost” vs. “non-roost” class), *L*_*reg*_ a regression loss for the predicted coordinates of the bounding boxes, and *λ* is a balancing parameter. The loss calculation and the optimization of the parameters are performed by splitting the training dataset into small batches. The training batch size is limited by memory constraints and is a hyper-parameter of the model. We evaluate our object detection models using the Mean Average Precision (mAP) at a 0.5 intersection over union (IoU), denoted as mAP@.5, as used in PASCAL VOC (Everingham *et al*., 2012). The IoU is a measure of similarity calculated between the ground truth bounding box and the predicted bounding box. If the IoU is greater than 0.5, the prediction is considered a true positive, otherwise it is considered a false positive. Mean average precision is a commonly used summary metric for detection models. First, average precision (AP) is calculated as the area under the curve of the precision-recall curve for each category, and then mAP is computed as the mean AP over all classes. Since ours is a single-class detection problem, mAP@.5 corresponds to the AP for bird roosts considering predictions with an IoU greater than 0.5 as true positives.

#### Tracking

We employ a simple greedy heuristic to associate single-frame detections into tracks (Ren, 2008). We start with high scoring detections and add unmatched detections in neighboring frames with high overlap. If a detection has enough overlap to be assigned to more than one track, we add it to the longest track. We apply the Kalman filter (Kalman, 1960) to smooth the roost tracks using a linear dynamical system for the bounding box center and radius. The linear system captures the dynamics of roost formation and expansion with parameters estimated from the ground truth annotations (e.g., the rate of expansion of roost bounding boxes). If no detection is matched in the neighboring frame due to missing or inaccurate single-frame detections, we use the bounding box predictions from the linear system. Lastly, we apply non-maximum suppression to remove duplicate roost tracks: if two tracks have enough roost detections with high overlap, we remove the shorter track. Refer to Cheng *et al.* (2020) for the tracking implementation details.

### 2.3 Dataset

We leverage the same radar scans and annotations with the same splits into training, validation, and testing sets as Cheng *et al.* (2020) does, but develop the dataset into the commonly adopted COCO format for computer vision datasets. After removing the scans with rendering errors, the three splits respectively have 53,266, 11,599, and 23,587 scans. These radar scans are manually annotated for prior ecological research (Laughlin *et al*., 2016) by different annotators. Each label records the position and radius of a circle that approximates the roost. Since the annotators have different annotation styles, Cheng *et al.* (2020) proposed a latent-variable model and an expectation-maximization algorithm (Dempster *et al*., 1977) to jointly learn a detection model and scaling factors specific to annotators. We adopt their learned factors to scale and standardize the annotations in the dataset. The training, validation, and test have 37,619, 5139, and 10,942 roost labels, respectively.

### 2.4 System Deployment

In the summer of 2020, we trained and deployed a version of our system to collect data about roosting behavior of swallows and martins using data from 12 radar stations in the Great Lakes region. The radar stations include KAPX, KBUF, KCLE, KDLH, KDTX, KGRB, KGRR, KIWX, KLOT, KMKX, KMQT, and KTYX; locations are shown in Figure 4. We use the deployed system to evaluate the effectiveness of an AI system for studying roosting behavior and to develop biological case studies, and as a baseline model for further improvements to the AI system.

**Figure 4:**
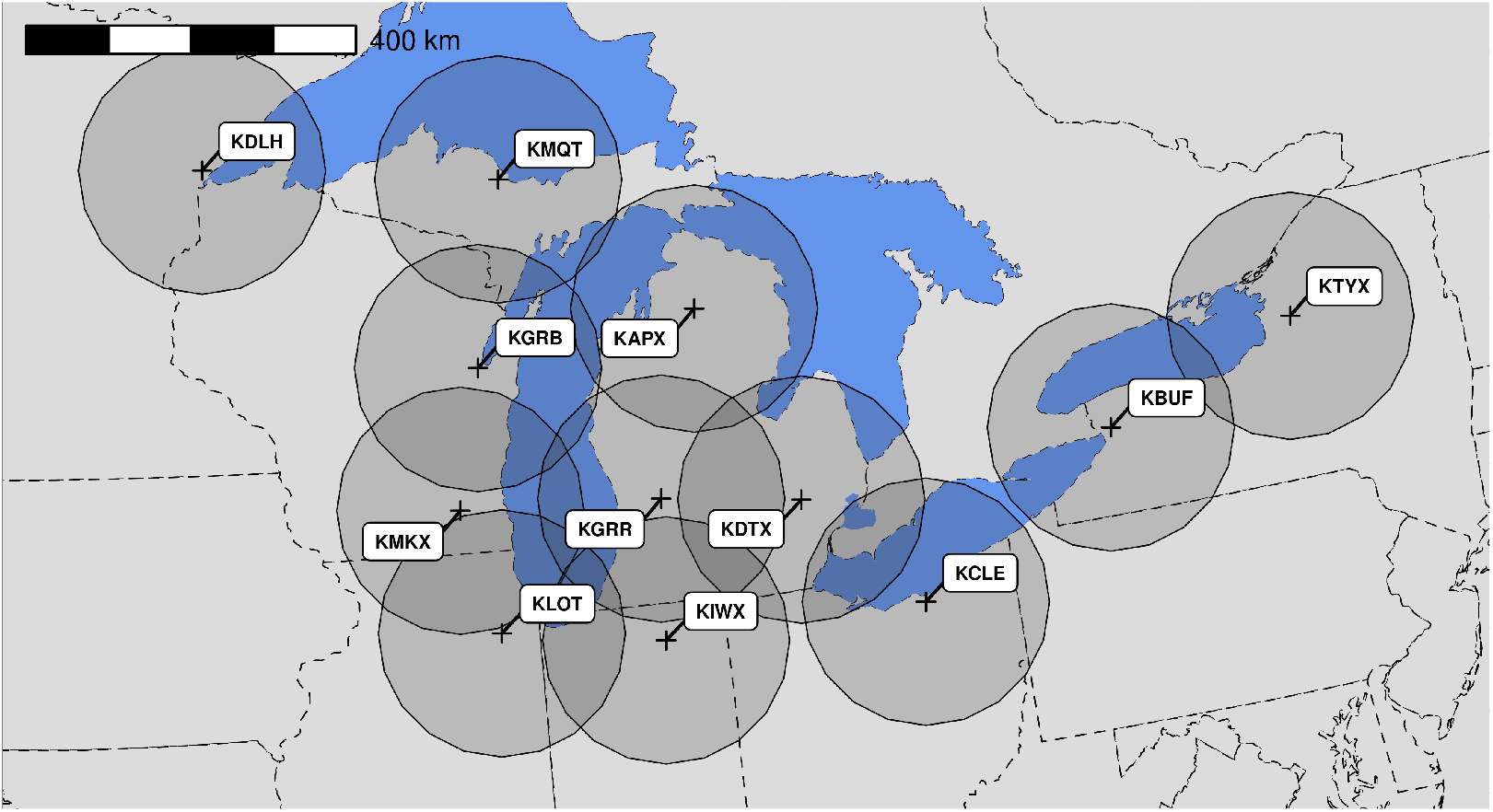
A map for 12 WSR-88D radar stations around the Great Lakes, which are a large group of five interconnected fresh-water lakes in the mid-east of North America, near the US-Canada border.

We downloaded scans from 30 minutes before to 90 minutes after local sunrise between June 1 and October 31 of the 21 years ranging from 2000 to 2020. Each station’s local sunrise times are obtained using the PyEphem (Rhodes, 2011) and pytz Python packages. In each 120-minute window, we set 41 reference times spaced 3 minutes apart; for each, we selected the scan closest to the reference time, or none if there were no scans within three minutes of the desired time. We downloaded and rendered each scan as a set of 600 × 600 arrays as described in Section 2.1, representing a 300 km × 300 km geographical region centered at the radar station.

We trained a Faster R-CNN detector with a ResNet101-FPN backbone pretrained on ImageNet classification and MS-COCO detection and 45 anchors ranging from 16 × 16 to 512 × 512. The detector takes three channels as input, for which we selected reflectivity at 0.5° and 1.5° and radial velocity at 0.5°. We scale the three-channel input to 1200 × 1200 to increase the spatial detail the network can resolve. We summarize the best training hyper-parameters in Appendix A. The deployed detector achieves a 46.98 mean average precision on the test dataset described in Section 2.3. During deployment, we kept the 100 top-scoring roost detections with predicted probability scores of at least 0.05 in each scan for further processing. We selected this low score threshold to ensure high recall, so that the predictions captured almost all roosts and could be screened in later processing stages to remove non-roosts.

The raw detections were assembled into tracks according to Section 2.2. We further post-processed tracks to remove rain and windfarms when possible, and to tentatively remove low-scoring tracks. We considered a detection as rain and removed it if the majority of pixels in its bounding box had copolar cross-correlation coefficient *ρ*_*HV*_ *>* 0.95, following the common rule of identifying rain (Dokter *et al*., 2018a). We removed detections corresponding to wind farms using known turbine locations from the U.S. Wind Turbine Database (Hoen *et al*., 2019). We built a web-based interface to display the remaining tracks together with the underlying radar imagery for further screening by humans. In the interface, any track with at least 2 detections, an average detection score of at least 0.15, and at least 1 detection with a score at least 0.5 was given the initial label of *roost* and were considered “high confidence” roosts; all other tracks were given the initial label of *non-roost* and were considered “low confidence” roosts. High-confidence tracks were displayed with full opacity for further screening; low-confidence tracks were displayed faintly (with low opacity) and could be ignored if they did not correspond to a true roost.

We developed a screening protocol to classify the predicted tracks into 7 categories. Each scan was screened by one of our three screeners (MD, YD, VS) following the proposed protocol: If more than half of a scan was filled with weather, all detections within this scan were removed from the analysis, and the whole scan was considered as missing data since we couldn’t have observed roosts if they had occurred. In other scans, clear roosts were labeled as *roost*. Roosts contaminated by weather, anomalous propagation, and unknown noise were labeled as *weather-roost, ap-roost*, and *unknown-noise-roost*, respectively. Some tracks corresponded to true roosts but had problems such as terminating early or late, drifting from the roost to another object, or including unreasonably large or small bounding boxes; these were labeled as *bad-track* to inform downstream analyses of issues with the track. These last five categories are suitable for ornithology research but with varying degrees of difficulty when analyzing the raw radar returns to estimate bird numbers. If the system detected multiple tracks that followed the same underlying roost, only one track was kept and others were labeled as *duplicate* tracks. Tracks that did not correspond to bird roosts at all were labeled as *non-roost*.

### 2.5 Detection Experiments

We conducted experiments to investigate the potential to improve the deployed system by incorporating additional input information into the detection model. We focused on the detection model independently from tracking because most errors in our initial system could be traced to detections errors that were difficult to correct with tracking alone. For example, when assembling single-frame detections into tracks, our tracking system also removes sporadic detections in isolated frames that do not behave like roosts; however, the improvement in performance after removing these false positives is negligible (∼0.2%).

We used a linear adaptor as described in Section 2.1 to augment the detection model to accept any number of channels, and systematically explored the use of all 15 modalities available (5 elevations × 3 legacy radar moments). Taking into account that the number of parameters of our model increases with the number of input channels, we performed experiments using different combinations of radar moments (i.e., reflectivity, velocity, and spectral width) and elevations (i.e., from 0.5° to 4.5°) to find the optimal set of inputs.

In addition to the information available in a single time frame of the radar data, we explored the use of additional temporal information by providing as inputs different combinations of radar data from previous and future consecutive frames. We included the consecutive frames as additional input channels to train our network (See Figure 2b). Our experiments used input data ranging from 3 to 45 channels (i.e., when using 3 consecutive frames with 3 radar moments at 5 elevations).

For completeness, we performed ablation tests changing one configuration at a time from our best model. The experiments conducted to examine the benefits of different design choices are summarized below.

- *Additional modalities*. The radar modalities are the 3 radar moments (i.e., reflectivity, velocity, and spectrum width), each one available at 0.5°, 1.5°, 2.5°, 3.5°, and 4.5° elevations. We used an adaptor between the input image and the first convolutional layer of the ‘backbone’. The adaptor linearly combines an arbitrary number of channels into 3 (the same number of channels required by the pretrained model). We ran tests with different combinations of radar moments and elevations (See Table 1).
- *Temporal information*. We tested using additional consecutive frames in combination with the frame *x*_*t*_ for which detections are predicted. We explored different combinations of up to 3 consecutive previous frames (up to *x*_*t−*3_) and up to 2 consecutive future frames (up to *x*_*t*+2_). All additional time frames were added to the input with the same number of channels (radar moments and elevations) used for *x*_*t*_ (See Table 2).
- *Pretraining*. We compared models with parameters initialized using ImageNet to models with further pretraining using MS-COCO (Lin *et al*., 2014), and to models trained with randomly initialized parameters.
- *Detector backbone*. We performed experiments using a ResNet50-FPN and a ResNet 101-FPN to check the performance of the detector with models of different capacity.
- *Input size*. Since the input size affects the amount of computation and memory needed by the network, we tested sizes from 800 × 800 to 1200 × 1200 to find the tradeoff between input size, which affects runtime memory and speed, and detection accuracy.
- *Anchors*. We performed tests with 3 different sets of anchor sizes for the region proposal network (RPN) of the Faster R-CNN. Because roost shapes are approximately isometric, we used a single 1:1 ratio for each size. The first used the default sizes from Faster R-CNN (5 sizes × 1 ratio = 5 anchors). In addition, we tested sets of 25 and 45 anchors; details are given in Appendix B.
- *Batch size*. We tested training batch sizes of 2, 4, and 8 samples.

**Table 1:** Additional modalities. Performance of the roost detector with different combinations of radar moments and elevations. Increasing the number of modalities from our baseline is not helpful. However, removing radial velocity at 0.5° and reflectivity at 1.5° harms performance. The mAP of the best model is shown in bold. A check mark (✓) indicates the included modalities in each experiment.

| Reflectivity |  |  |  |  | Velocity |  |  |  |  | Spectrum width |  |  |  |  | mAP@.5 |
| --- | --- | --- | --- | --- | --- | --- | --- | --- | --- | --- | --- | --- | --- | --- | --- |
| 0.5 | 1.5 | 2.5 | 3.5 | 4.5 | 0.5 | 1.5 | 2.5 | 3.5 | 4.5 | 0.5 | 1.5 | 2.5 | 3.5 | 4.5 |  |
| ✓ | ✓ | ✓ | ✓ | ✓ | ✓ | ✓ | ✓ | ✓ | ✓ | ✓ | ✓ | ✓ | ✓ | ✓ | 54.39±1.7 |
| ✓ | ✓ | ✓ | ✓ |  | ✓ | ✓ | ✓ | ✓ |  | ✓ | ✓ | ✓ | ✓ |  | 54.06±0.8 |
| ✓ | ✓ | ✓ |  |  | ✓ | ✓ | ✓ |  |  | ✓ | ✓ | ✓ |  |  | 54.22±1.0 |
| ✓ | ✓ |  |  |  | ✓ | ✓ |  |  |  | ✓ | ✓ |  |  |  | 54.34±1.2 |
| ✓ | ✓ |  |  |  | ✓ |  |  |  |  | ✓ |  |  |  |  | 54.11±1.2 |
| ✓ | ✓ |  |  |  | ✓ |  |  |  |  |  |  |  |  |  | **54.71±1.0** |
| ✓ | ✓ |  |  |  |  |  |  |  |  |  |  |  |  |  | 53.90±1.1 |
| ✓ |  |  |  |  |  |  |  |  |  |  |  |  |  |  | 53.83±1.3 |

**Table 2:**
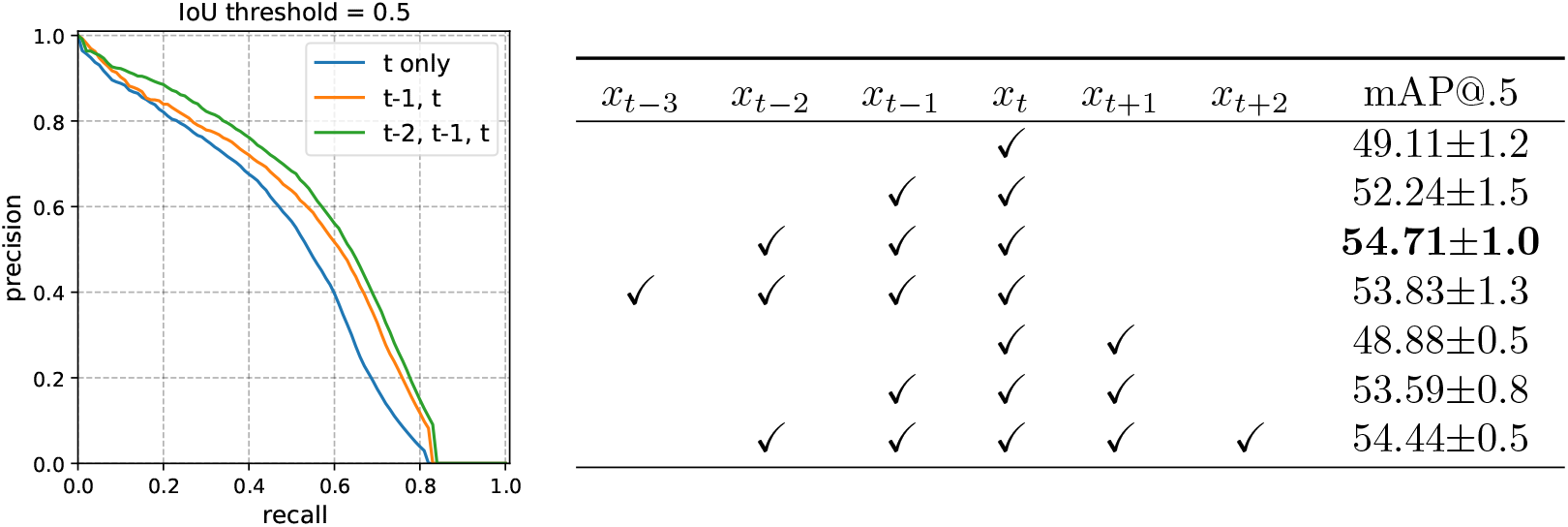
Temporal information improves detection performance. **(Right)** Performance on target frame (*x*_*t*_) with different combinations of previous and future frames. Most combinations improve performance. However, using future frames alone is not helpful. The mAP of the best model, which uses *x*_*t*_, *x*_*t−*1_, and *x*_*t−*2_, is shown in bold. Check marks (✓) indicate the time frames included in each experiment. **(Left)** Precision-recall curves when using a single frame (*t* only), two consecutive frames (*t* − 1, *t*), and the best model with three consecutive frames (*t* − 2, *t* − 1, *t*).

### 2.6 Biology Case Studies

To demonstrate the types of ecological questions that could be answered with our machine learning pipeline, we selected a dense subset of tracks found in the Walpole Island First Nation Reserve, in southwestern Ontario, Canada. This region is within the ranges of both the KDTX and KCLE stations, which could result in the same roost being detected twice. For the purposes of this paper, we selected tracks captured only by KCLE. After manually screening the detections to remove contaminants, as described above, we used the bounding boxes estimated by the model to extract the raw reflectivity factor values from Level II radar data in polar coordinates (the location of each voxel or sampling volume is defined by its range and azimuth) across all elevation angles of each full scan.

The number of birds in each bounding box (in each roost) can be estimated using the approach established by Chilson *et al.* (2012). The method can only be implemented for tracks that were labelled as clear roosts (593 tracks from our case study region), since it relies on the assumptions that each bounding box only contains biological scatterers and that the birds are uniformly distributed within each voxel, here defined by the region illuminated by the radar beam for each range gate. We convert equivalent reflectivity factor (*Z*_*e*_), originally in dBZ, to linear scale (mm^6^ mm^*−*3^), and then transform it into reflectivity (*η*), which we interpret as the density of scatterers in the atmosphere (in cm^2^ km^*−*3^). We can then multiply the reflectivity measurement at each azimuth and interval by the theoretical volume sampled by the radar at that range. Finally, we divide the result by the specific radar cross section (RCS) of the bird species assumed to be found in the roost, thus obtaining an estimate of the number of birds.

To obtain conservative counts, we used Purple Martins as our benchmark radar cross section, since they are the largest species of Hirundinidae found in North America. The radar cross section of Purple Martins can be obtained from their mass - 51g, see (Dunning, 2008) - adopting the relationship between mass and RCS proposed by Horton *et al.* (2019a): log(RCS) = 0.699 × log(mass). Bank Swallows (*Riparia riparia*), possibly the smallest species to participate in such aggregations, would yield counts approximately times higher, since their mass is 13g according to Dunning (2008). The volume sampled by the radar is assumed to be shaped like a truncated cone with axis aligned with the antenna’s peak power axis, cut by two parallel planes at each range gate. For data before the Super Resolution upgrade, the cone’s apex angle was assumed to be 1°. After the 2007-2008 upgrade, we assumed an elliptical cone of 1° vertical beam width and 0.5° horizontal beam width (Torres, 2007).

We extracted the number of birds from all sweeps available in each radar scan. In a post-processing stage, we filtered the sweeps that had height lower than 5000m within the 150km radius of each station. The sweeps were then grouped in bins of 1° interval according to their elevation angle, and we calculated the mean count within each bin. Finally, we summed the estimates from each elevation bin to get the bird count of each roost. This procedure is needed to avoid double counting, because we assumed a 1° vertical beamwidth.

With the extracted bird count of each roost detection, we then explored phenological trends at the spatial scale of this roost, which consistently occurs from 2000 to 2020. Data was considered as missing on days when scans had more than 50% weather contamination, intense anomalous propagation, or when the sampling window was shorter than 100 minutes. Days without detections were considered as true zeros. To calculate the daily bird count within the roost for each day, we derived the maximum number of birds for each roost track. We then fit a generalized additive model (GAM) to each roost-year to model the roosting activity throughout a roosting season. We constructed GAMs with daily bird counts as the response variable and ordinal date as the independent variable with the smoothing parameter *k* set to 5 using a quasi-Poisson distribution. We used this model construction to predict counts throughout the season and selected the 50% passage date, i.e., the first date in which the cumulative predicted counts exceeded half of the yearly total, as our phenology estimate for that year.

## 3 Results

We will first report results about the performance of the detection model, and then report results about the system deployment and biological case studies. All models were compared using mean average precision (mAP) at 0.5 IoU. We found that performance did not change significantly after training for more than 40k iterations. Therefore, we ran each experiment 3 times for 50k iterations and evaluated the trained models at 40k, 45k, and 50k iterations. For each experiment, we report the average mAP and standard deviation over those 9 evaluations.

Our deployed model, described in Section 2.4, achieved a mAP of 47.0%, compared to 45.5% for the best performing detector in Cheng *et al.* (2020). Both models are based on a Faster R-CNN and use a single timeframe image as input. In the following sections we describe the improvements over the deployed model.

### 3.1 Overall performance

The best performing model overall included the adaptor layer and used as input 3 consecutive time frames, *x*_*t−*2_, *x*_*t−*1_, and *x*_*t*_, with three input channels per frame: reflectivity at 0.5°, reflectivity at 1.5°, and radial velocity at 0.5°. These 9 total input channels were rendered at size 1100× 1100 as the inputs to a Faster R-CNN detection model with linear adaptor and a pretrained ResNet101-FPN backbone, then trained for more than 40k iterations with a batch size of 4 samples. The model had mAP of 54.71% on the test set. All other settings were the same as the deployed model (Section 2.4). As shown in Figure 5, this model detects most roosts with few false positives even when the radar images contain rain (bottom of the radar scans in Figure 5) and non-roost biological scatterers or anomalous propagation (center of the radar scans). See Appendix C for more qualitative results.

**Figure 5:**
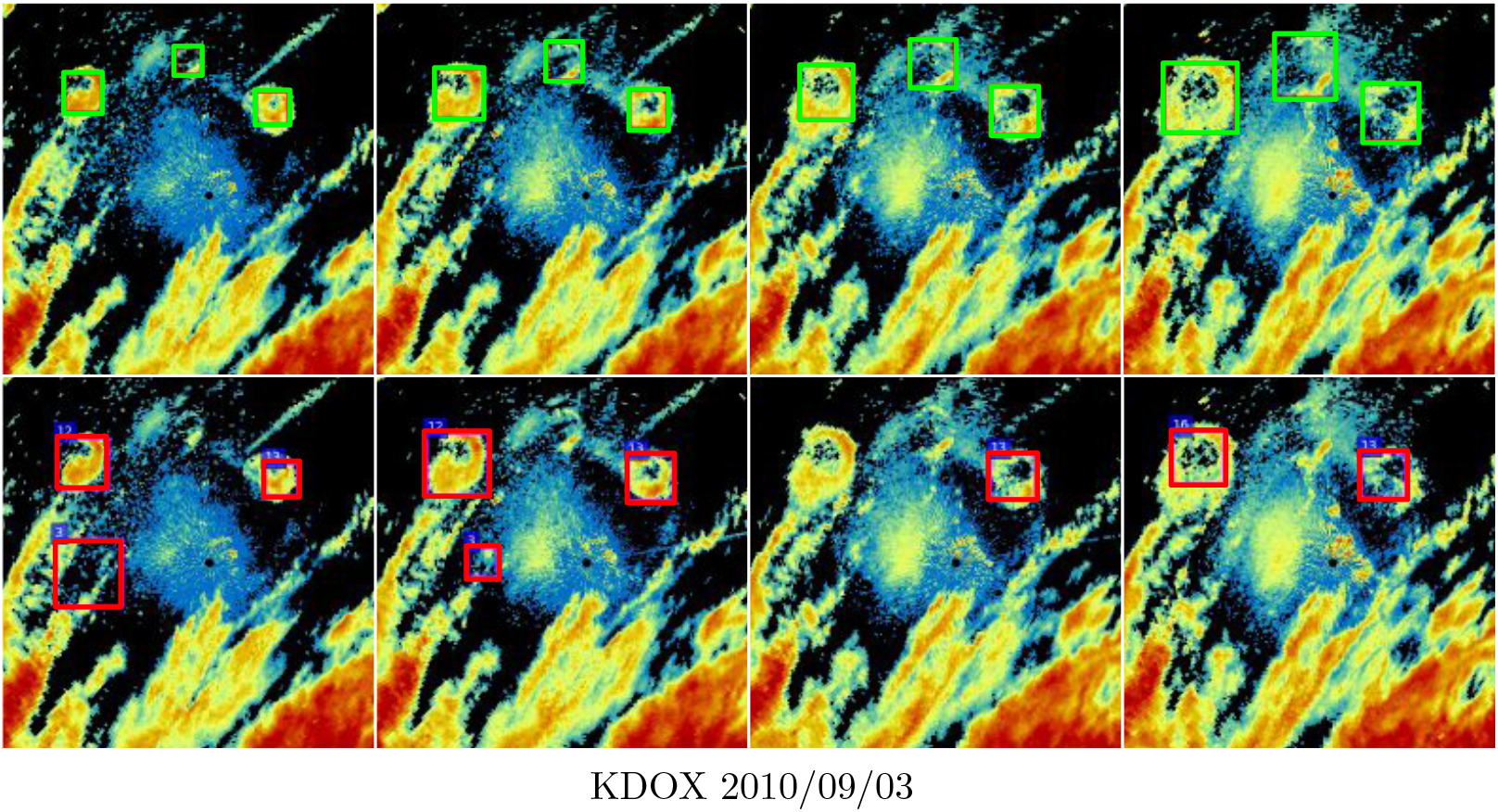
Qualitative Results. Detections on 4 consecutive frames in the test set. We show human annotations with green color in the top row and predictions of our roost detector with red color in the bottom row. KDOX radar station on 2010/09/03.

### 3.2 Detection experiments

This section presents the results of ablation experiments to evaluate the impact of removing individual model components from the best overall model as described in Section 3.1.

- *Additional modalities*. Table 1 shows results with different combinations of radar moments at different elevation angles used as input to the detector. Overall, adding additional modalities did not improve performance compared to using only reflectivity at 0.5°, reflectivity at 1.5°, and radial velocity at 0.5°. For instance, the 3-channel model had mAP of 54.71%, the model that used all radar moments at all elevations had mAP of 54.39%. This result is not completely unexpected since communal bird roosts are mainly found in low elevations. Using fewer than 3 input channels, however, harms performance: models with 2 and 1 channels achieve mAP of 53.90% and 53.83%, respectively.
- *Temporal information*. Table 2 shows results with different combinations of consecutive time frames used as input. Adding temporal information to the model helped considerably (5.6% mAP increase). These extra time frames likely allow the model to detect qualitative differences in the movement behavior of roosts in radar imagery compared to weather and other biological scatterers. For example, rain patterns are highly variable and often include small regions that appear ring-shaped like roosts, however, the movement dynamics are very different: rain usually moves in an approximately straight trajectory and deforms slowly, while roosts diverge from a single point. Adding more than two previous frames did not improve the model’s performance: specifically, the model with 3 previous frames had mAP of 53.83%, while the model with 2 previous frames had mAP of 54.71%. Including previous frames did not just increase mAP, which is an aggregate performance metric; it led to higher precision at all prediction thresholds (different levels of recall in the precision-recall curve; Table 2, left).

Surprisingly, adding future time frames made almost no difference in model performance. Adding a single future time frame gave mAP of 48.88% compared to mAP of 49.11% without future frames. We hypothesize that roosts are more challenging to detect in their later stages when they dissolve and fade from view. Thus, using information from previous time frames helps during these later stages, but a roost in a future time frame is expected to be even more challenging to detect and not add helpful information to the model.

An interesting finding is that the adaptor layer improved performance even without additional input channels: the deployed model had mAP of 46.98%, and the model with the same inputs plus an adaptor layer had mAP of 49.11% (first row of Table 2). The adaptor layer may provide a useful way to transform the input images to be most suitable for use with the filters in the early layers of the pre-trained CNN.

- *Pretraining*. Table 3 compares models with different pretraining schems. The model pretrained with classification and detection datasets (ImageNet+MS-COCO) outperformed (54.71%) the model pretrained only with ImageNet (53.28%). To prevent the model trained with randomly initialized weights from diverging, we had to significantly decrease the learning rate (from 10^*−*3^ to 10^*−*7^). To compensate for the slower learning rate, we increased the number of training iterations by ∼35×. After these changes, we could train the model but it still had significantly worse performance (39.72%) than the two pretrained models.

**Table 3:**
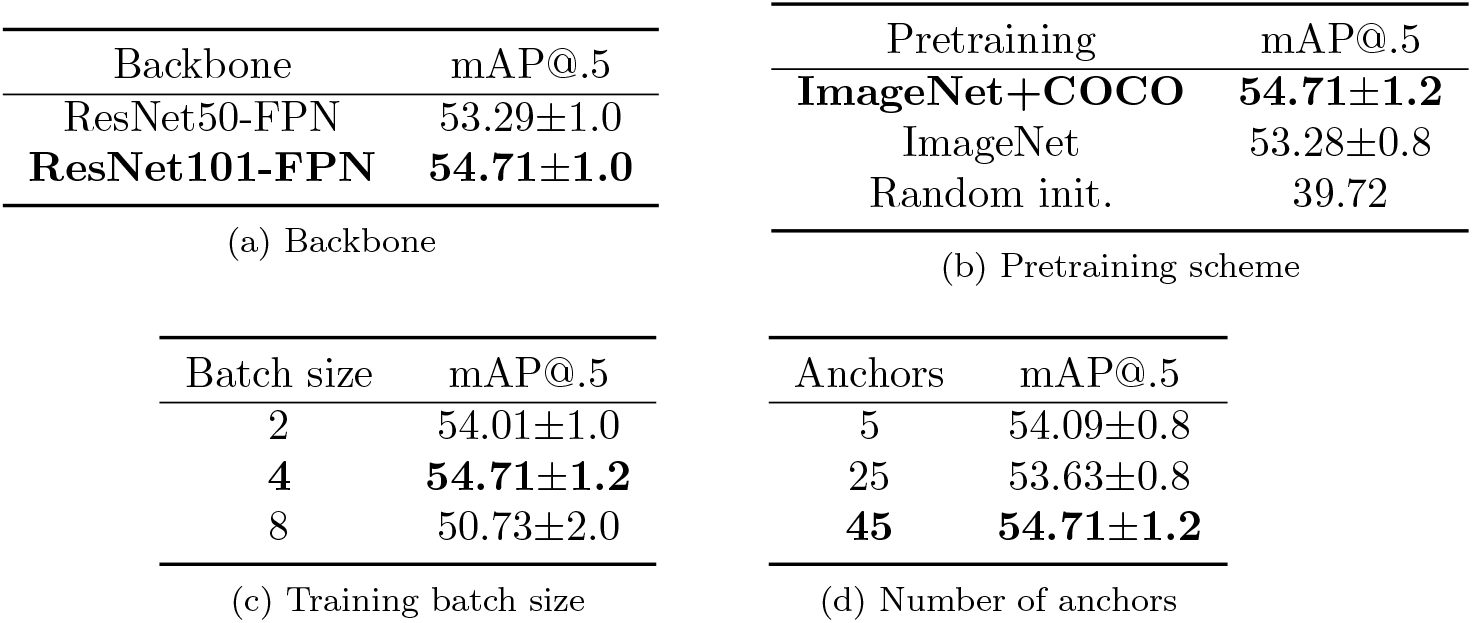
Performance with different design choices. Performance with varying backbone, pretraining scheme, batch size, and number of anchors. These experiments use input size 1200 1200. The best results use a batch size of 4 samples, 45 anchors, and a ResNet101-FPN backbone. The parameter settings used in the overall best performing model are shown in bold.
- *Detector backbone*. The detector with the higher capacity ResNet101-FPN (54.71%) outperformed the detector with a ResNet50-FPN backbone (53.29%) without overfitting the training data.
- *Input size*. Using an input size of 1200 × 1200 gave comparable results to 1100 × 1100. We decided to use 1100× 1100 since the running time and memory requirements were lower. Table 4 shows mAP, floating point operations (FLOPS), and memory usage with different input sizes.

**Table 4:**
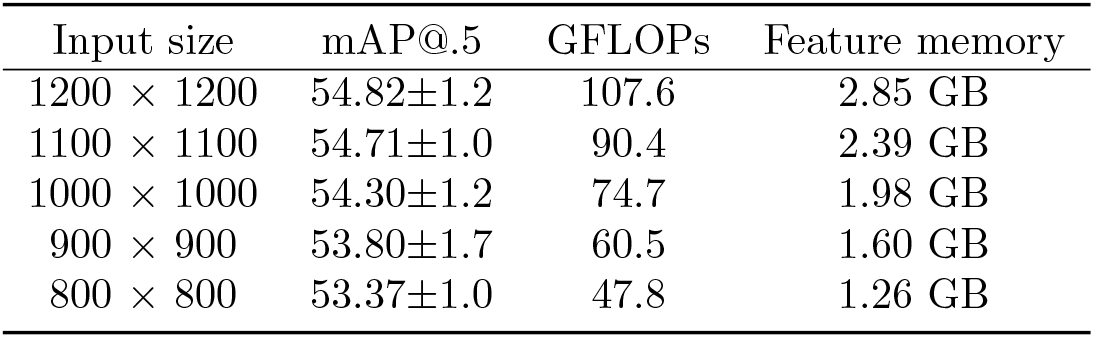
Radar image input size. Performance with different input sizes. An input size of 1200 × 1200 gives comparable results to an input size of 1100 ×1100, but uses less resources. Gigaflops and feature memory (RAM required to store the output of intermediate layers) are calculated for a single-image batch.
- *Anchors*. Table 3 shows results with different number of anchors in the Faster R-CNN region proposal network. The model with 45 anchors was best with performance of 54.71%, compared to 53.63% with 25 anchors and 54.09% with 5 anchors.
- *Batch size*. Due to memory constraints, we could only evaluate batch sizes of up to 8 samples. A batch size of 4 led to the best performance (54.71%), and increasing the batch size to 8 decreased detection performance by ∼ 3.3% (Table 3). In some cases, small mini-batch sizes provide more up-to-date gradient calculations, which yields more stable and reliable training (Masters & Luschi, 2018).

| Input size | mAP@.5 | GFLOPs | Feature memory |
| --- | --- | --- | --- |
| $1200 \times 1200$ | 54.82±1.2 | 107.6 | 2.85 GB |
| $1100 \times 1100$ | 54.71±1.0 | 90.4 | 2.39 GB |
| $1000 \times 1000$ | 54.30±1.2 | 74.7 | 1.98 GB |
| $900 \times 900$ | 53.80±1.7 | 60.5 | 1.60 GB |
| $800 \times 800$ | 53.37±1.0 | 47.8 | 1.26 GB |

### 3.3 System Deployment

Table 5 summarizes the statistics for the deployed system described in Section 2.4. There were six station-years for which data were not available or could not be rendered: the year 2000 for KDLH and KGRB; and the years 2001-2004 for KTYX. For the remaining station-years, 612,786 scans were successfully rendered in total. Our system predicted 31,313 high-confidence tracks assembled from 140,036 detections. Another 230,088 tracks with 372,594 detections were designated as low confidence before screening.

**Table 5:**
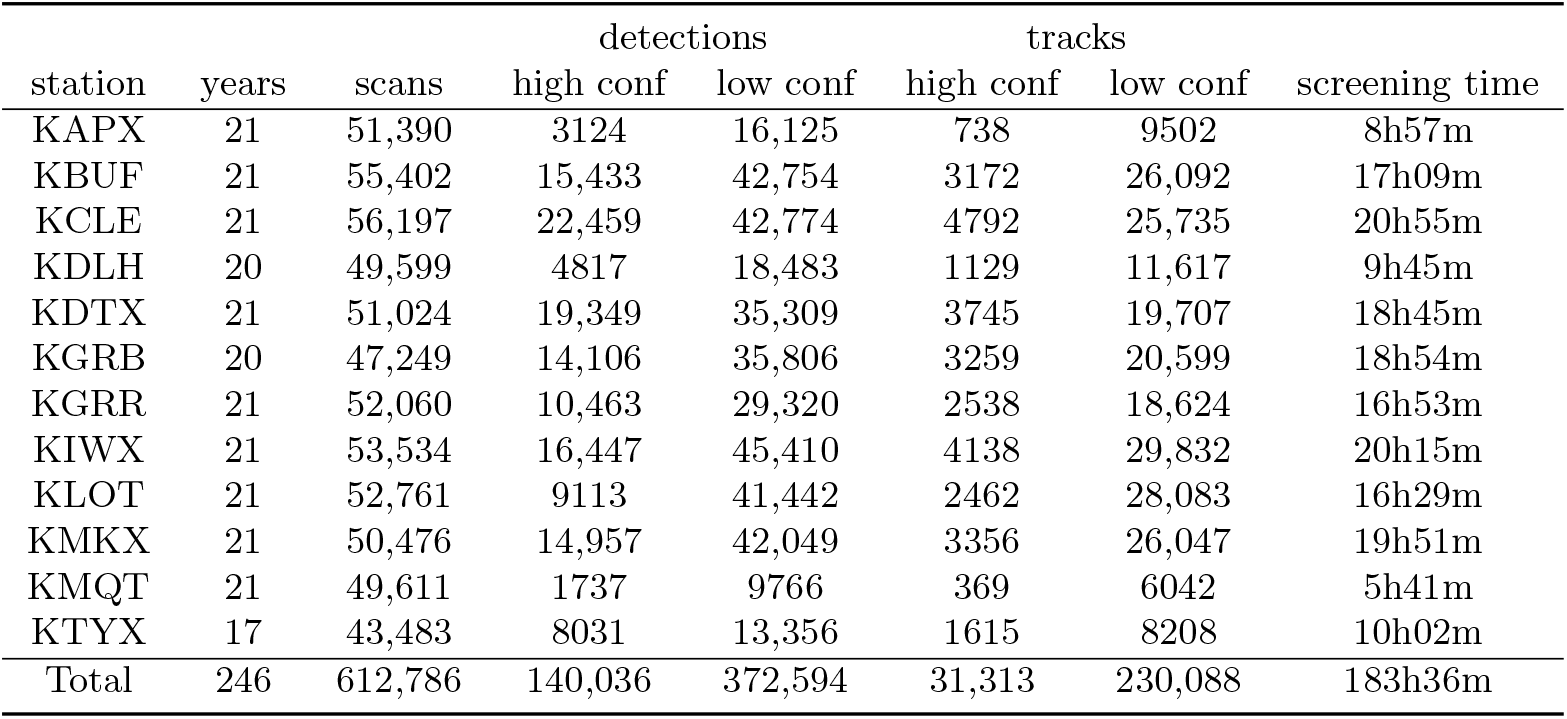
Statistics for rendered scans, system predictions, and time needed to screen the predictions. System-predicted detections and tracks are either of high or low confidence and displayed as *roost* or *non-roost* by default in the screening interface; see Section 2.4 for details. The bounding boxes are shown with high and low opacity, respectively, to ease screening.

Table 6 shows the statistics for the screened data. From the 612,786 scans, our annotation pipeline identified 13,860 clean roost tracks, 477 tracks that were contaminated by weather, 100 that were contaminated by anomalous propagation, and 1191 that contained unknown noise. These four categories together produced 15,628 roost tracks of 64,620 detections that are useful for ornithology research. Among the low confidence predictions, only 3603 tracks (8266 detections) out of 230,088 tracks (372,594 detections) were marked as one of the four roost categories; a large number of *non-roost* s did not require any action from the screener. The screening amounts to 183.6 annotator hours and an average of 1.08 second per scan. If the annotators were to find roosts, draw bounding Backbone mAP@.5 ResNet50-FPN 53.29±1.0 boxes for all roosts, and assemble them into tracks completely manually, the annotation process would be significantly more time-consuming and error-prone. Our AI system reduces human labor to screening system predictions and enables tractable annotation to obtain high quality roost data.

**Table 6:** Statistics for screened data. Entries show numbers of detections/tracks assigned each of the 7 labels: clean roost (cln), weather contaminated roost (wth), anomalous propagation contaminated roost (ap), unknown noise contaminated roost (nse), duplicate (dup), bad track (bad), and non-roost track (non). The first four categories can be used for aeroecology research.

| station | cln | wth | ap | nse | dup | bad | non |
| --- | --- | --- | --- | --- | --- | --- | --- |
|  | number of detections / number of tracks |  |  |  |  |  |  |
| KAPX | 1040/311 | 14/1 | 18/4 | 22/6 | 5/3 | 32/7 | 18118/9908 |
| KBUF | 6669/1628 | 384/79 | 72/11 | 1129/156 | 90/30 | 334/45 | 49509/27315 |
| KCLE | 12863/3017 | 800/151 | 53/12 | 903/155 | 46/16 | 585/82 | 49983/27094 |
| KDLH | 2249/679 | 52/15 | 54/13 | 136/30 | 13/6 | 49/10 | 20747/11993 |
| KDTX | 9568/1990 | 729/117 | 81/12 | 578/95 | 74/28 | 659/75 | 42969/21135 |
| KGRB | 3616/998 | 94/20 | 51/12 | 1364/292 | 15/6 | 725/116 | 44047/22414 |
| KGRR | 4827/1340 | 68/13 | 31/9 | 572/138 | 23/7 | 346/44 | 33916/19611 |
| KIWX | 7017/1890 | 118/25 | 98/21 | 632/146 | 4/1 | 449/65 | 53539/31822 |
| KLOT | 1841/610 | 39/11 | 1/1 | 186/42 | 2/2 | 50/9 | 48436/29870 |
| KMKX | 1911/510 | 90/19 | 10/2 | 520/91 | 5/2 | 95/14 | 54375/28765 |
| KMQT | 9/7 | 0/0 | 0/0 | 0/0 | 0/0 | 0/0 | 11494/6404 |
| KTYX | 3752/880 | 146/26 | 11/3 | 202/40 | 24/11 | 1116/147 | 16136/8716 |
| Total | 55362/13860 | 2534/477 | 480/100 | 6244/1191 | 301/112 | 4440/614 | 443269/245047 |

### 3.4 Biology Case Studies

Figure 6a shows the locations of the first detection in each of the 702 tracks, colored by year, in the study area at Walpole Island around Lake Saint Claire. Martins and swallows gathered in this region to roost every year from 2000 to 2021, with an average of approximately 33 days of detections per year. The yearly maximum number of days when birds were detected occurred in 2019, when birds congregated around the lake on 42 days between July 23rd and September 8th.

**Figure 6:**
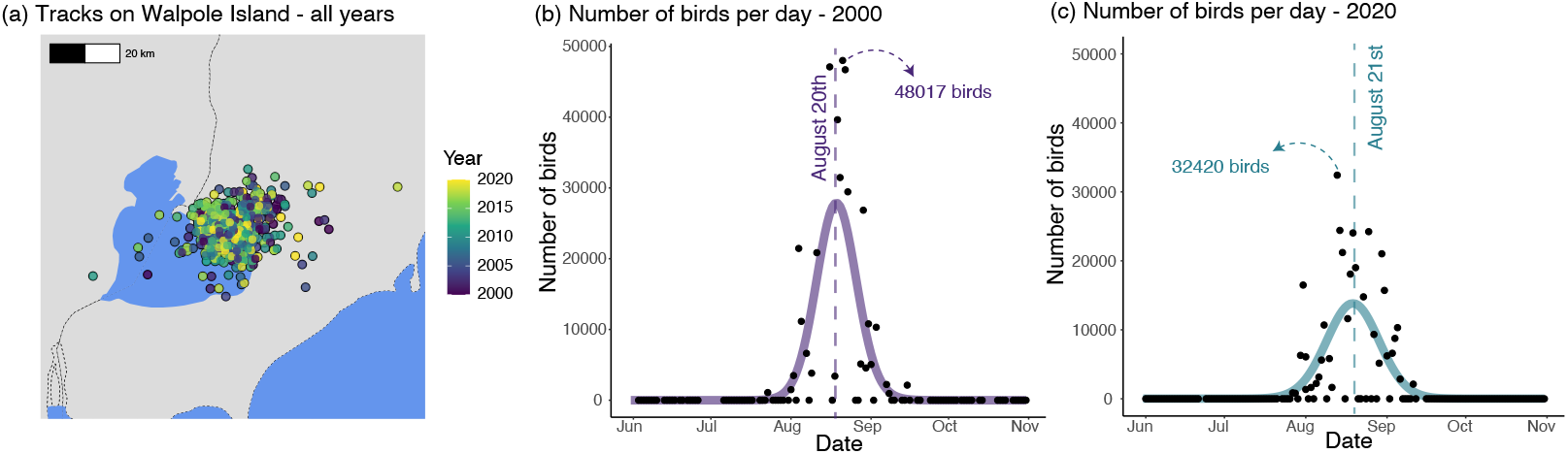
Case Study roost at Walpole Island, Ontario, Canada. (a) Map where points, colored by year, represent the first detection of each track captured by the KCLE radar station. (b) Raw bird counts (represented as points) and the GAM prediction (continuous line) of roosting activity from June to October, 2000. (c) Raw bird counts (represented as points) and the GAM prediction (continuous line) of roosting activity from June to October, 2020.

The peak timing (50% passage date) for roosting activity in the study area was August 20th in 2000, and August 21st in 2020 (Figure 6 b-c). The average peak number of birds per year detected in the region throughout our study period was 73,925 birds (SD detections tracks = 37,743). The year when the roost received the highest number of birds occurred on 2010, when our counts reached 156,864 on August 8th. In contrast, the year with lowest peak count was 2013, when we detected at most 23,949 birds on August 9th.

## 4 Discussion

NEXRAD data represent one of the largest potential ecological datasets, spanning nearly three decades. In this paper we developed an AI-assisted roost detection system to unlock significant and critical insights from this spatial and temporally dense dataset, and applied the system to collect research-grade data on roosts in the Great Lakes region of the US.

### Sources of false positives

Future research can focus on improving an AI-system through a careful error analysis. Figure 7b shows the distribution of top scoring false positives (detection with a score of 0.5 or higher) for the temporal detector. Localization errors (IoU*>*0) account for nearly 44% of the false positives, while detections on background regions (IoU=0), frequently due to weather, static structures, or anomalous propagation, account for the remaining 56%. Figure 7a shows the precision-recall curve by varying the overlap threshold at which a detection is considered a true positive. The mAP improves from 56.3% to 66.4% when the overlap threshold is 0.4 compared to 0.5. However, some roosts are still entirely missed by the model (see false negative sources below). See examples of localization errors in Figure 8a-c, missed roosts in Figure 8d, and detections on background regions in Figure 8e.

**Figure 7:**
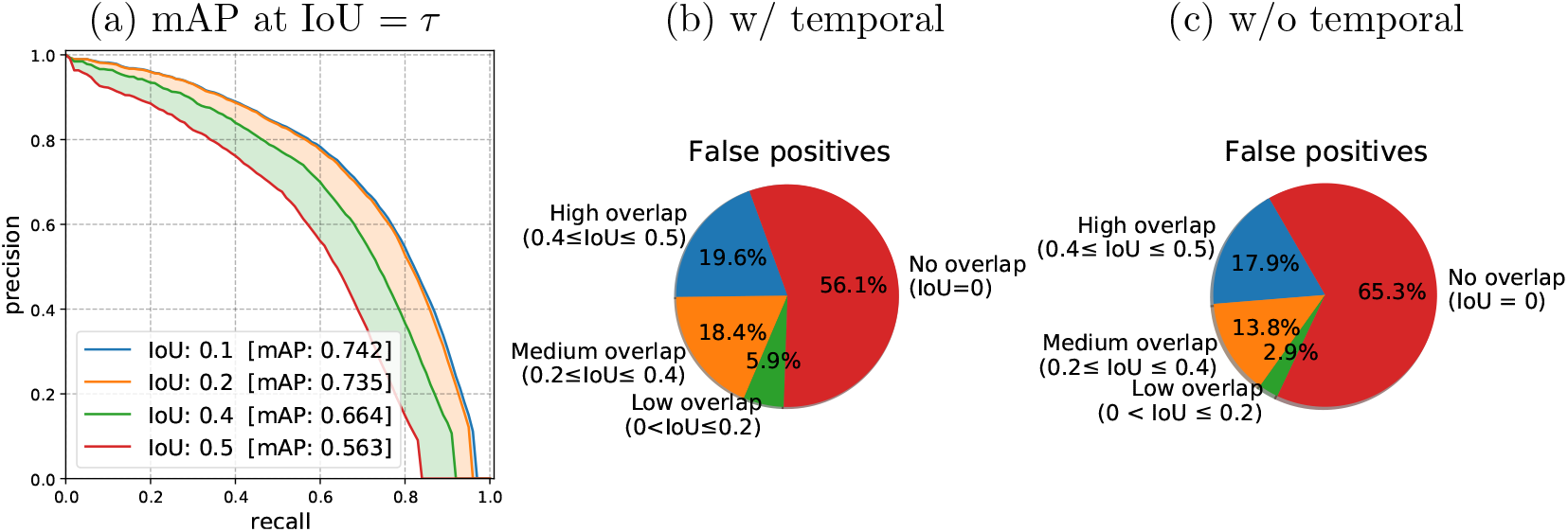
**(a)** Precision-recall curves at different IoU thresholds (higher values reflect a stricter requirement). The mAP improves from 56.3% to 66.4% when a threshold IoU*>*0.4 is used compared to IoU*>*0.5 suggesting that a large number of top-scoring detections are “near misses”. **(b)** The distribution of top scoring false positives for the model with temporal information. Localization errors (IoU*>*0) account for nearly 44% of the data, while detections on background regions, frequently due to weather, account for the remaining 56.1%. **(c)** The distribution of top scoring false positives without using temporal information shows a greater fraction of false positives on the background regions.

**Figure 8:**
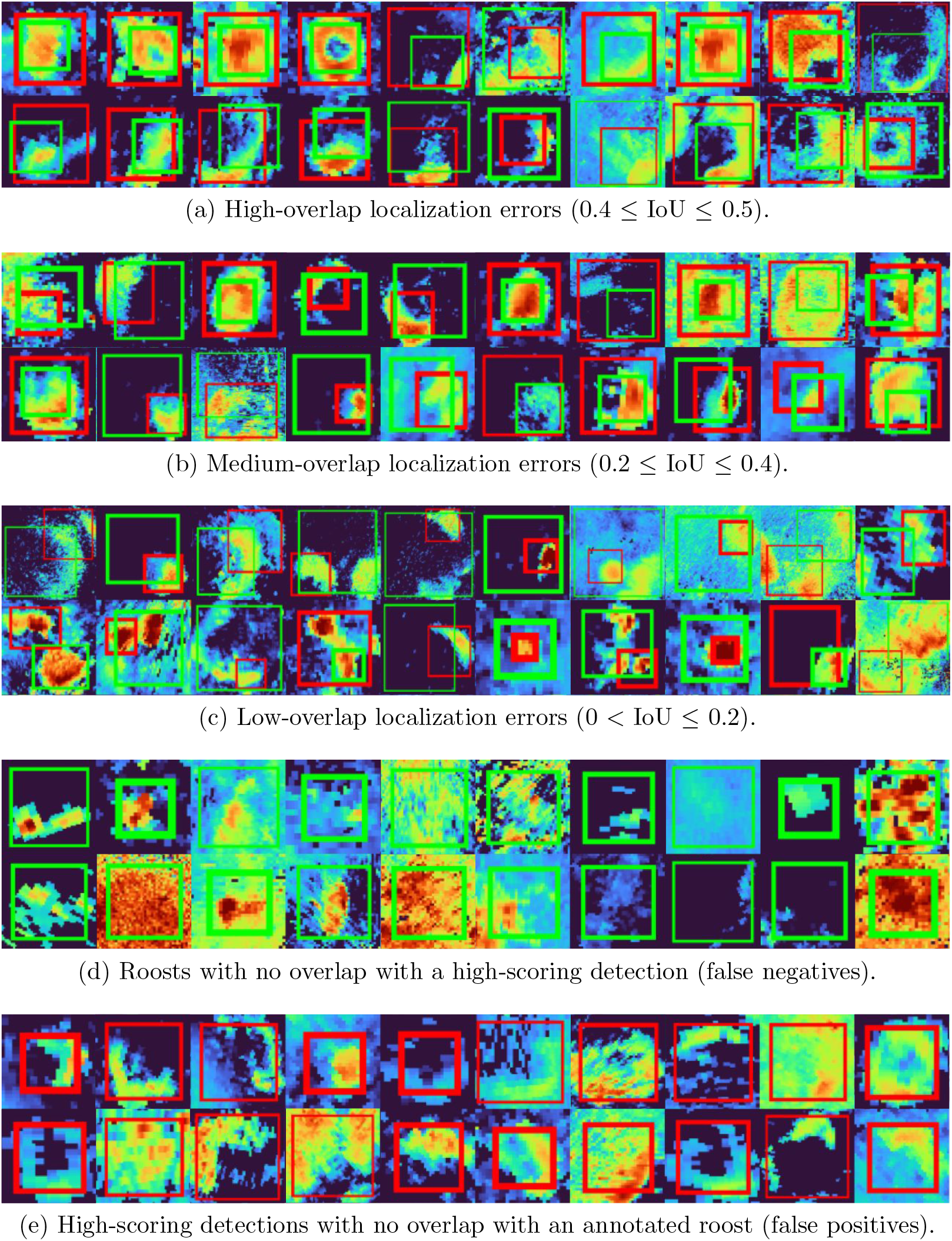
Detection errors. A detection is considered false positive if the overlap (IoU) between the annotation and prediction bounding boxes is ≤ 0.5. **(a-c)** Detection with different overlap values below the threshold. **(d)** Human-annotated roosts that the model does not detect. **(e)** High-scoring detections on rain show a similar morphology as roosts in a single frame, but the temporal model is able to reduce many of these false positives. We show human annotations in green and predictions by the model in red.

### Temporal information reduces false positives

A significant technical advance was to incorporate temporal information from past frames to capture roost dynamics. When viewed in a size-constrained window rain can take many different shapes, including the appearance of a ring-shaped roost (see examples in Figure 8e). However, the movement of weather is different from roosts—rain often moves in an straight trajectory, while roosts diverge from a point. Temporal information reduces false positives due to small patches of rain with a roost-like appearance (Figure 9a). Similarly, static structures such as wind farms and buildings appear as small “blobs” of high-reflectivity and are often confused as small roosts by the single-frame detector. Using multiple frames reduces these false positives due to the differences in dynamics (Figure 9b). The overall distribution of top-scoring false positives on background was reduced from 65.3% to 56.1% with the addition of temporal information (Compare Figure 7b and Figure 7c).

**Figure 9:**
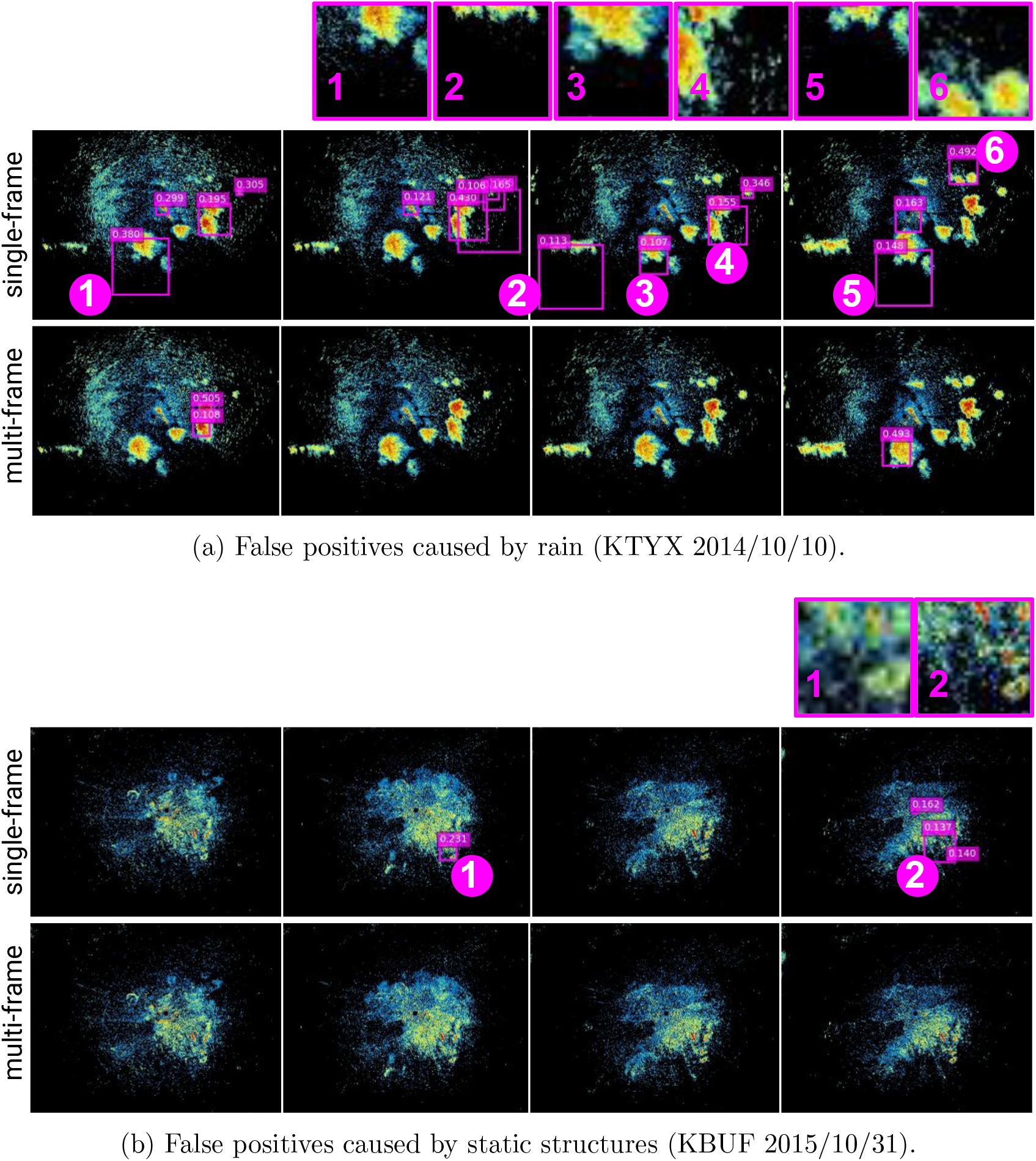
Temporal information reduces false positives. Detection results by our detector (bottom row) compared with a detector trained with a single time frame *x*_*t*_ (top row). **(a)** KTYX radar station on 2014/10/10 10:54:32–11:10:22 UTC. **(b)** KBUF radar station on 2015/10/31 11:34:48–12:04:12 UTC. There are no bird roosts in the frames shown, so all detections are false positives.

### Sources of false negatives

We found several sources of false negatives, which are roosts that were annotated by researchers who originally created our training data set (Laughlin *et al*., 2016) but missed by our system. The first source is radar scans that contained roosts together with large amounts of other noise such as anomalous propagation (AP), ground clutter, or other biological scatterers. Figure 10a shows an example of three consecutive frames with a single annotated roost (green) amidst AP and ground clutter with reflectivity values so high that the model is unable to discriminate the roost from the background. In this case, the human labeler likely relied on contextual information such as the presence of roosts in the same location across many frames and days (see more examples of noisy roosts in Figure 8d). Also note that the model successfully detects roosts in radar scans with AP when the reflectivity values are not so high (see examples in Figure 5 and Figure C.1a in the Appendix C).

**Figure 10:**
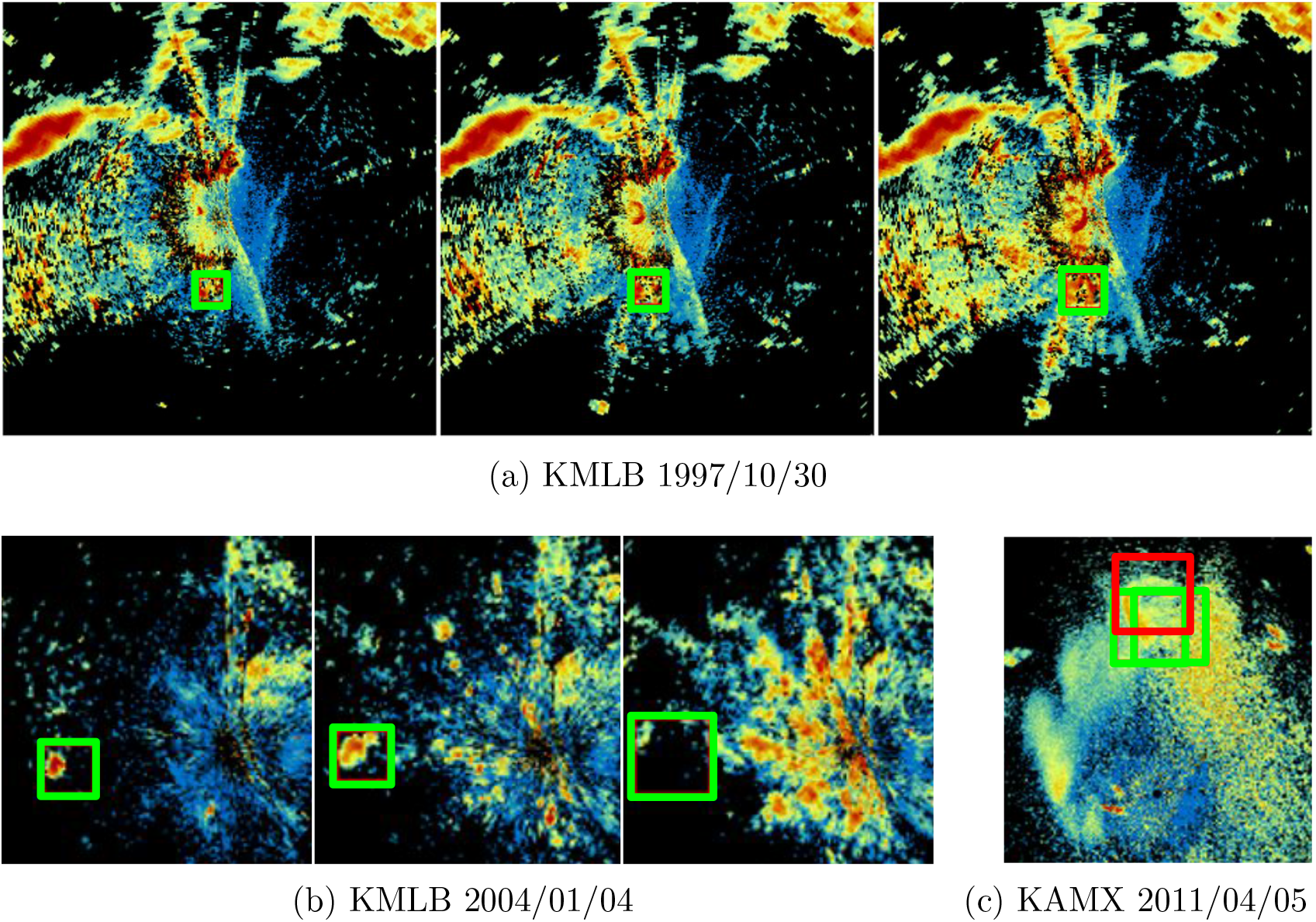
False negative sources. **(a)** Three consecutive radar scans with a large amount of anomalous propagation and other biology. **(b)** Three consecutive scans with a roost that lacks a well-defined ring shape. **(c)** Single scan with two roosts overlapping – the model only detects one. We show human annotations in green and predictions if produced by the model in red. Images correspond to the reflectivity at 0.5° elevation.

Another source of false negatives is roosts that lacked the prototypical ring shape. Figure 10b shows annotations for a non-ring-shaped roost in three consecutive frames. These are encountered rarely in the dataset and the morphology can easily be confused with weather and other biology (see more examples of roosts without ring shape in Figure 8d). Figure C.1c shows an example where the model *is* able to detect a roost without the usual ring shape; however, there is a higher correspondence between consecutive frames and a more consistent diverging pattern compared to the example in Figure 10b. Detection of non-ring-shaped roosts could be complicated by the fact that annotators were required to use a circle to annotate roosts, and often adopted different labeling styles, especially for how to label a roost that was not ring shaped (Cheng *et al*., 2020).

One more source of false negatives is bird roosts that appear too close to each other in the radar scan. As shown in Figure 10c, the model detects only one of the two annotated roosts. Again, the human annotators may have used complex contextual information to interpret these overlapping patterns as two different roosts.

### Future research for AI-assisted roost detection

There are a number of future directions to improve an AI-assisted system for roost measurement. Iterative refinements to an existing model can often improve performance significantly over time, especially when driven by a careful error analysis. Adding new sources of information, including new input features, training signals, or training labels, together with data curation efforts to improve the quality of training data, are among the most promising ways to improve performance. In the specific context of our work, adding the results of human-screened predictions back into the model as training data may provide a significant boost in model performance to scale our model to larger geographical regions with less human effort.

Future research can incorporate additional sources of information related to the contextual cues that humans use to detect roosts and discriminate them from other patterns in radar images. One such cue is the persistence of roosts in the same or similar locations across days and years, which may allow humans to confidently detect roosts in noisy radar scans including high-reflectivity AP, weather, or other biological scatterers. A model could be given input features from radar scans in previous days or years, similar to the way we added within-day temporal information in this paper; or, it could be given historical *detections* of a simpler roost detection model (Zhou *et al*., 2020). We found that rain is a persistent source of false positives, even though humans have a relatively easy time distinguishing rain from roosts using the full context of a radar image sequence, and AI models can successfully discriminate rain from broad-scale bird migration (Lin *et al*., 2019). There may be a mismatch between the “local” scale of an object detection model and the broader scope of contextual information needed to discriminate rain. A possible remedy is to train a multi-head model to jointly perform roost detection together with another task such as rain segmentation (Kirillov *et al*., 2018; Li *et al*., 2022; Shen *et al*., 2021; He *et al*., 2017), or provide the predictions of a rain segmentation model (Lin *et al*., 2019) as input to the roost detection model.

Ultimately, the goal is to collect roost measurements from a very large but finite set of images. Future research should focus on effective ways to combine human effort, computational effort, AI training, and statistical estimation to achieve the desired scientific outcomes. For example, what is the tolerance of measurements of swallow phenology or population declines to an AI system with a certain performance level? Is human screening of outputs required? What are the most effective strategies for interleaving human annotation and model training to analyze a very large image data set? An interesting research direction is to pose the scientific question (e.g., how many roosts) as a statistical estimation problem and to consider statistical estimators that can give high confidence bounds after examining only a subset of the images (Meng *et al*., 2021).

### Broader directions for biological recognition

There are several broader directions in the recognition of biological patterns in radar data that can be informed by our work. Our roost-detection model can serve as a starting point for AI models to detect and track related taxa-specific biological phenomena in radar data, including bat roosts (Stepanian & Wainwright, 2018), roosts of non-swallow bird species (Van Den Broeke, 2019), and mayfly hatches (Stepanian *et al*., 2020). Of these, bat roosts are most similar to swallow roosts, and may require the least adaptation to the model. As a proof of concept, we deploy our best model found in Section 3.2 on bat roosts around the KEWX radar station in Texas. We process scans from 90 minutes before local sunset to 150 minutes after in June 2012. We observe that our system developed using bird roost data performs reasonably well at detecting and tracking bats without any customization (Figure 11).

**Figure 11:**
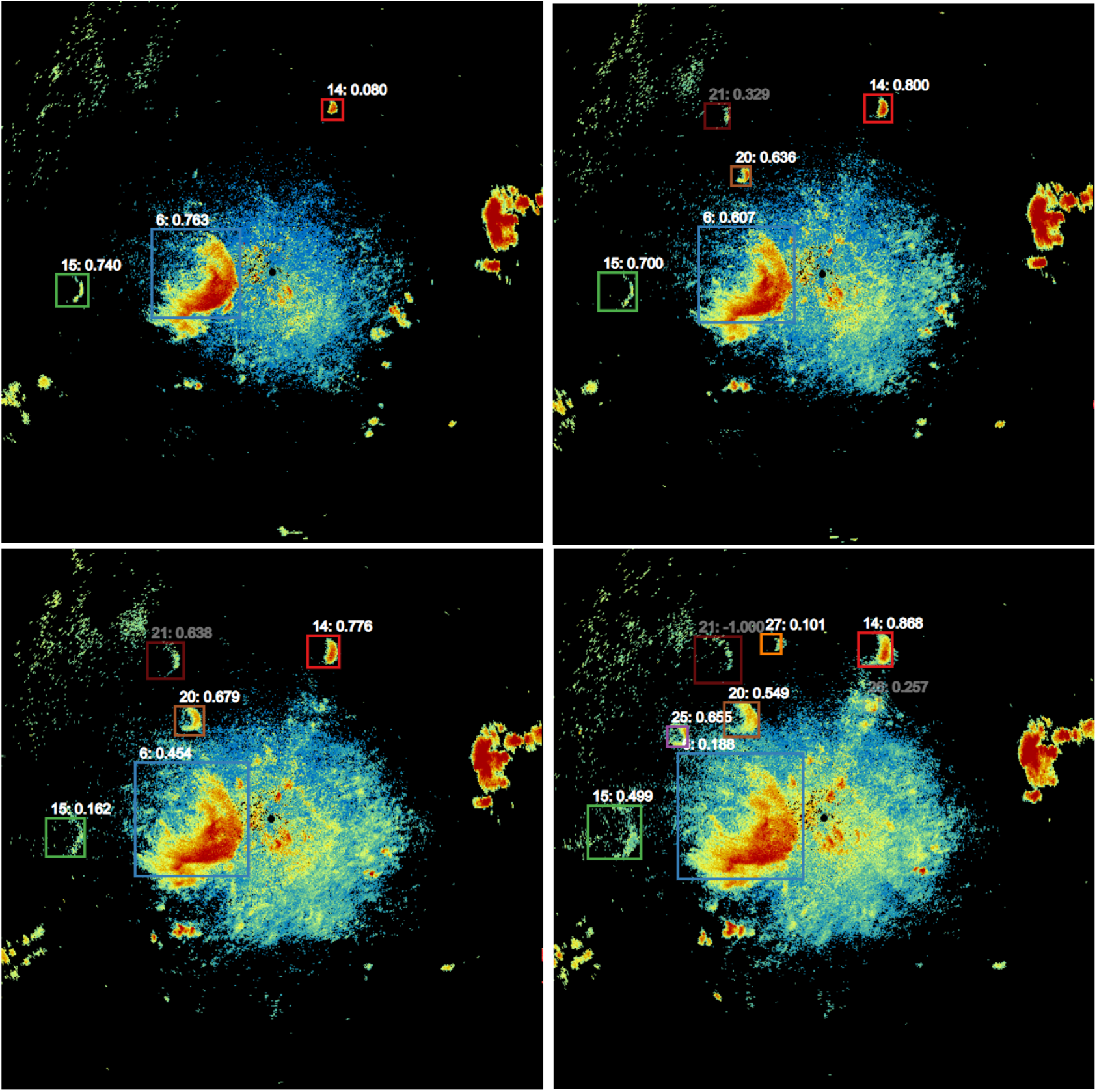
System detection and tracking of bat roosts from the KEWX radar station on 2012/6/30. The scans are temporally consecutive in the order of top-left, top-right, bottom-left, and bottom-right.

Roosts of other bird species including robins, blackbirds, starlings, and waterfowl are also visible on radar (Russell *et al*., 1998), but are often less obvious for humans to discern and usually lack the distinct “expanding ring” pattern of swallow roosts, probably due to differences in roost emergence behavior. Mayfly hatches have a distinct and rather different appearance than bird roosts. The automatic detection of these phenomena is an interesting frontier for AI methods in radar aeroecology. Investigating whether it is possible to distinguish among roosts of different swallow species, e.g., Purple Martins and Tree Swallows, from fine-grained radar characteristics is an interesting open question that could have important biological implications. Finally, extending these models to radar networks outside the US could provide information to track bird species beyond national borders.

## Acknowledgments

This material is based upon work supported by the US National Science Foundation under Grant Nos. 2017554 to Colorado State University, 1749833 and 2017756 to the University of Massachusetts Amherst, and 2017582 to the University of Oklahoma.

## Author contributions

DS and SM conceived the study. WZ, ZC and SM created training data sets. WZ and ZC wrote code for model training, evaluation, and deployment. GP, WZ, ZC, and SM developed neural network models. GP developed final models and ran experiments. GP and SM analyzed results. WZ deployed model on Great Lakes data. DS wrote code to visualize and screen results. MB and YD designed screening protocol. MB, YD, and VS executed screening protocol. MB and YD analyzed data for biology case study. GP, DS, WZ, SM, MB, YD, and ZC wrote the manuscript. All authors contributed critically to the drafts and gave final approval for publication.

## A Deployed Hyper-parameters

This section summarizes the training hyper-parameters used to get our deployed detector. We show the specific sizes and ratios of the anchors in Appendix B.

**Table A.1:** Baseline training hyper-parameters.

| Training hyper-parameters |  |
| --- | --- |
| Backbone | ResNet101-FPN |
| Pretraining | ImageNet+COCO |
| Data augmentation | None |
| Input size | 1200 × 1200 |
| Input channels | 3 |
| Batch size | 4 |
| Learning rate | 1e-3 |
| Anchors | 45 |
| Training iterations | 50000 |

## B Anchor sizes

This section presents the anchor configurations for the region proposal network (RPN) used in our experiments. We get the best performance using set 3 of anchors.

**Table B.2:**
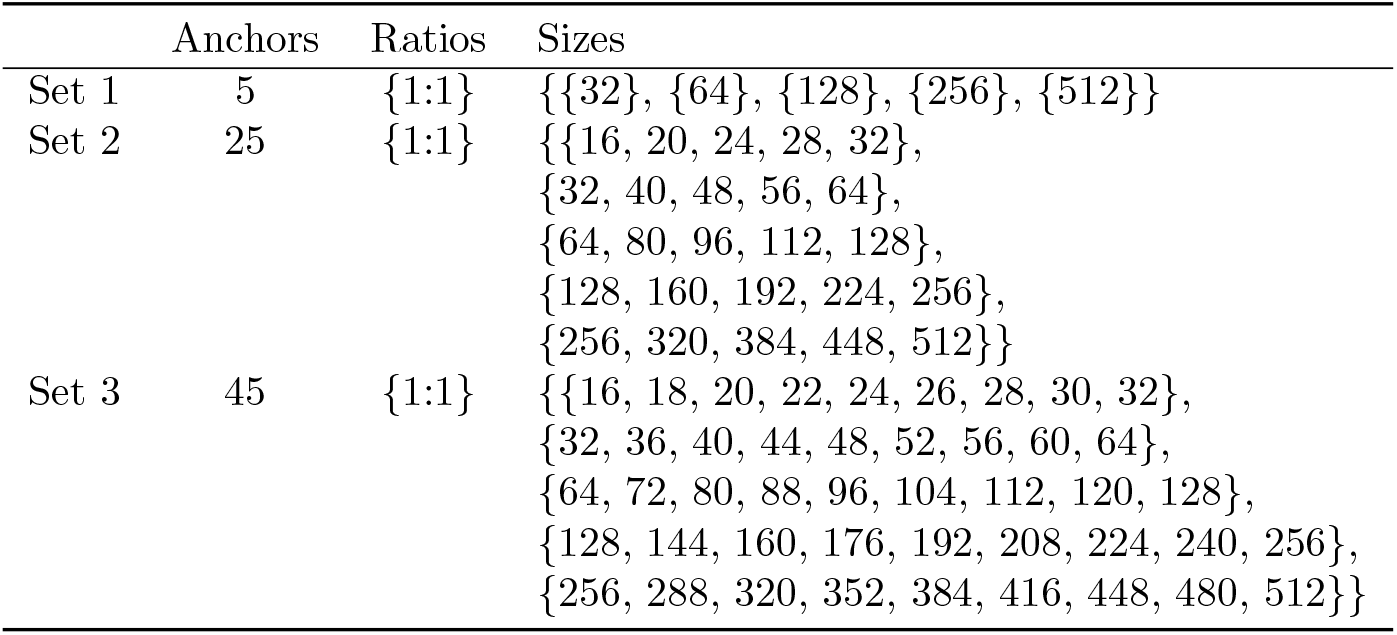
Baseline training hyper-parameters.

## C Additional qualitative results in the test set

**Figure C.1:**
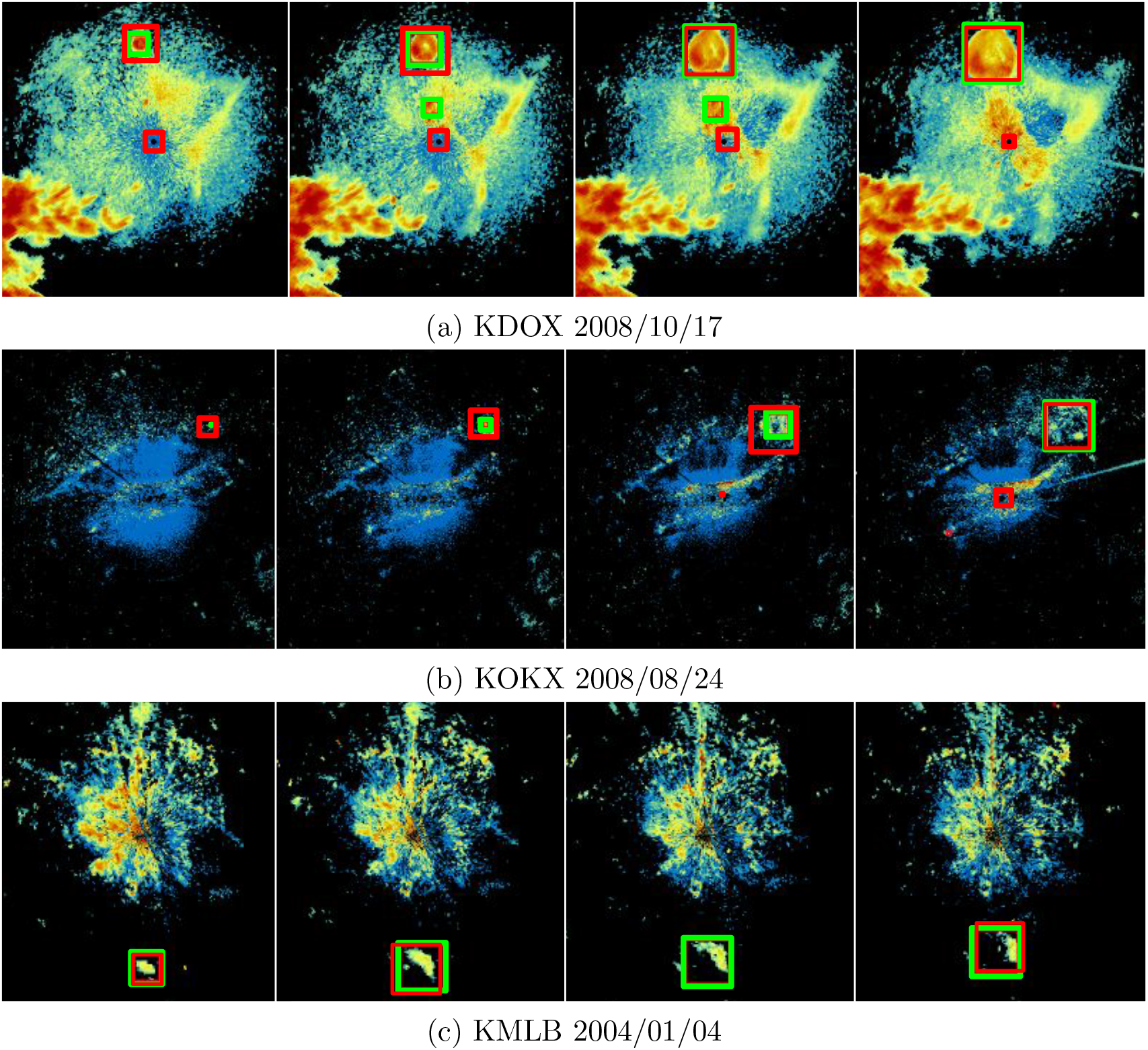
Qualitative Results. Detections on 4 consecutive frames in the test set on different stations. We show human annotations in green color and predictions in red.

## Footnotes

1 Each size value correspond to the square root of the area in pixels.

